# Molecular complexity of the major urinary protein system of the Norway rat, *Rattus norvegicus*

**DOI:** 10.1101/478362

**Authors:** Guadalupe Gómez-Baena, Stuart D. Armstrong, Josiah O. Halstead, Mark Prescott, Sarah A. Roberts, Lynn McLean, Jonathan M. Mudge, Jane L. Hurst, Robert J. Beynon

## Abstract

Major urinary proteins (MUP) are the major component of the urinary protein fraction in house mice (*Mus* spp.) and rats (*Rattus* spp.). The structure, polymorphism and functions of these lipocalins have been well described in the western European house mouse (*Mus musculus domesticus*), clarifying their role in semiochemical communication. The complexity of these roles in the mouse raises the question of similar functions in other rodents, including the Norway rat, *Rattus norvegicu*s. Norway rats express MUPs in urine but information about specific MUP isoform sequences and functions is limited. In this study, we present a detailed molecular characterization of the MUP proteoforms expressed in the urine of two laboratory strains, Wistar Han and Brown Norway, and wild caught animals, using a combination of manual gene annotation, intact protein mass spectrometry and bottom-up mass spectrometry-based proteomic approaches. Detailed sequencing of the proteins reveals a less complex pattern of primary sequence polymorphism than the mouse. However, unlike the mouse, rat MUPs exhibit added complexity in the form of post-translational modifications including phosphorylation and exoproteolytic trimming of specific isoforms. The possibility that urinary MUPs may have different roles in rat chemical communication than those they play in the house mouse is also discussed.

## INTRODUCTION

Physiological production of substantial protein in the urine is well known in both rats and house mice [1, 2]. The protein fraction is dominated by 18-19 kDa, eight stranded beta-barrel lipocalins known as major urinary proteins (MUPs, also named as α2u-globulins when first identified in rats [2, 3]). Urinary MUPs are a heterogeneous mixture of multiple isoforms that are very similar in mass and isoelectric point [4–6]. The functions of MUPs have largely been studied in the western European house mouse (*Mus musculus domesticus*) where they play critical roles in olfactory communication. First, they act as carriers for low molecular weight pheromones and other constituents, delaying their release from urinary scent marks [7–9]. MUP polymorphism also provides an identity signal for individual and kin recognition [10–14] and may play a role in species recognition [6]. Finally, MUPs act as pheromones in their own right [14–17]. In particular, darcin (MUP20, MGI nomenclature; http://www.informatics.jax.org/) has a number of unique properties, including a highly specialized role as a male sex pheromone that also induces competition between males. This protein binds most strongly the abundant volatile male pheromone (S)-2-(sec-butyl)-4,5-dihydrothiazole in mouse urine [16–19]. The pheromonal properties of darcin are retained in the recombinant protein showing that it acts as a pheromone in the absence of bound ligands [16, 17].

The structure and functions of MUPs in the house mouse are well established and serve to emphasise the significantly lower degree of understanding of the MUP system in rats, which differ in social organization from house mice [20]. Evidence is emerging that rat MUPs are likely to be important in male sexual and/or competitive communication, with urinary MUP output appearing around male puberty and increasing with a surge in testosterone levels [21, 22]. Male rats that are preferred by females express a greater amount of urinary MUP, and female rats are attracted to spend time near the high molecular weight fraction of male urine that contains rat MUPs and other urinary proteins [23]. Females also spend longer sniffing glass rods painted with castrated male urine if recombinant MUPs are added to the urine at normal physiological concentration [22]. Exposure to recombinant MUPs stimulates increased expression of the immediate early gene *c-fos* in the accessory olfactory bulbs of females and in brain areas known to be involved in pheromone-induced sexual behaviours [22]. However, the specific functions played by rat MUPs in sexual and/or intrasexual competitive communication have yet to be addressed, and it is not known whether different MUP isoforms play different roles as among mice.

While urinary MUP expression has been well characterized in mice and the urinary protein pattern can largely be reconciled with genome-level evidence [24–26], comparable information about the isoforms of MUPs expressed in rats is limited. There are no studies that have provided a deep analysis of the MUP protein complement in rat urine, a necessary step that precedes characterization of the function of the individual isoforms in communication. Although the rat genome sequence was first published in 2004 [27], gene annotation has lagged behind that of the mouse genome and it is more difficult to connect proteins observed in rat urine to the cognate coding sequences predicted from the genome sequence. Furthermore, phenotyping of individual urinary MUPs isoforms in rats has previously been based largely on 1D/2D-SDS-PAGE, or isoelectric focusing (with or without prior purification) [2, 3, 21, 22, 28–32]. Neither PAGE nor isoelectric focusing alone provides adequate resolution for the highly heterogeneous mixture of MUPs isoforms. By contrast, intact mass analysis by electrospray ionization (ESI-MS), complemented with mass spectrometry based protein sequencing, has proved a valuable tool for the characterization of the urinary MUP profiles in different species and strains [4–6, 25]. There have been some studies that make use of analytical mass spectrometry to study rat urinary proteome but none of these have addressed the issue of complexity and isoform phenotyping.

To gain a similar level of understanding on rat MUPs that exists for the house mouse, we have performed a manual annotation of the MUP genome cluster in the latest assemblies of the rat genome and completed phenotypic urinary profiling of male and female individuals from the laboratory strains Brown Norway and Wistar Han, and from some wild caught individuals. This strategy has allowed the detailed characterization of individual isoforms, reconciling genomic information and protein data, and has provided new insight into the post-translational modifications undergone by the rat MUP family, including phosphorylation and exoproteolysis.

## RESULTS AND DISCUSSION

### MUP genome cluster analysis

The first iteration of the sequenced rat genome was published in 2004 [33]. Since then, several assemblies have been released. We performed manual annotation of the rat MUP cluster on rat chromosome 5 using the genome assemblies RGSC_3.4 (v4) (December 2004), Rnor_5.0 (v5) (March 2012) and Rnor_6.0 (v6) (July 2014) from the Rat Genome Sequencing Consortium (Figure 1). Manual annotation revealed ten genes in v4 (numbered to maintain the nomenclature utilized by Logan and colleagues [26]) and eight in v5 and v6 (named A-H), along with several pseudo-genes (twelve in v4 and ten in v5 and v6).

**Figure 1.**
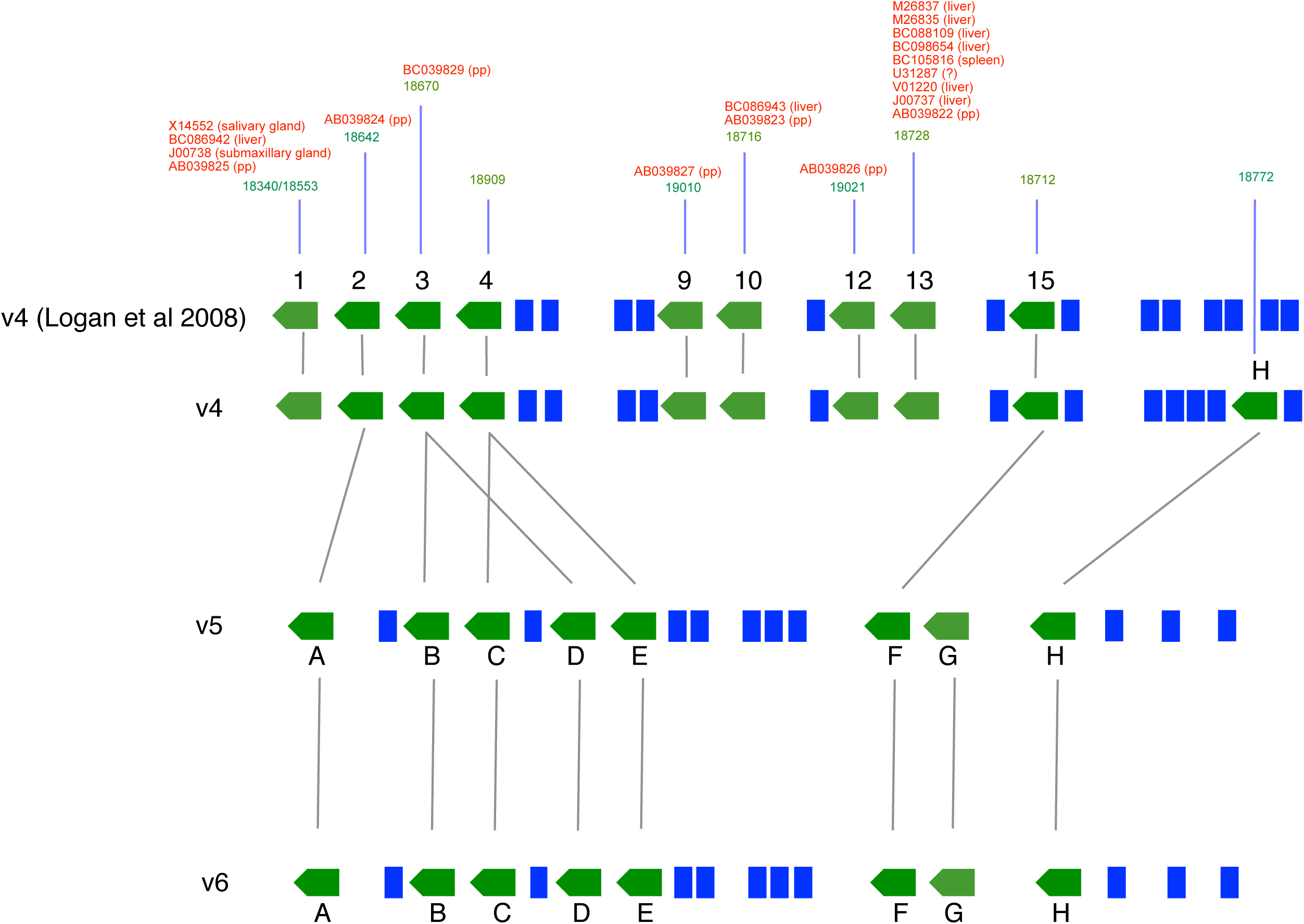
The MUP gene cluster of the rat. Three iterations of the rat MUP gene cluster have been produced in different genome assemblies. Besides, we show annotation reported by Logan et al [26] on v4. The gene identities between the assemblies are indicated by grey lines. Protein coding genes are green arrows and pseudogenes are blue boxes. In green, above each protein coding gene is the predicted mature mass for the protein in Da (corrected for signal peptide cleavage and a single disulfide bond formation). In red, transcriptional supporting information already available in the literature [43, 54, 58, 67], with the tissue of origin in brackets (pp: preputial gland).

The annotation at v4 accords well with that previously reported [26] (Figure 1) but with the addition of a single protein coding gene (gene H). Six out the 10 protein coding loci have transcriptional evidence, although only three in hepatic tissues (genes 1, 10, 13, yielding mature predicted masses of 18340, 18716 and 18728 Da respectively).

In v5 and v6, several genes (genes 1, 9, 10, 12 and 13 from v4) are removed compared to the v4 annotation, including two genes for which protein-level evidence had been obtained in urine; gene 13 (18728 Da) and gene 1 (18340 Da) [22, 34–36]. The fact that these putative genes (genes 1, 9, 10, 12 and 13 from v4) have transcriptional support indicates that these are genuine protein coding genes. Assembly v5 and v6 define a duplication of genes 3 and 4 that previously had single instances in v4. Additionally, genes F and G in v5 and v6 are incompletely covered in the genome sequence. Ultimately, the incomplete nature of the genome sequence across the rat MUP cluster compromises the ability to produce fully comprehensive gene annotations at this time. By contrast with the genome project of the C57BL/6 mouse, which utilized a hierarchical mapping and clone-based sequencing strategy, the rat genome sequences were generated almost entirely through whole-genome shotgun sequencing. We may anticipate that the highly duplicative nature of the MUP locus presents particular challenges to this strategy, including the assembly of DNA sequences into a correct genomic region. In fact, we cannot assume that the v5/v6 assemblies are necessarily of better quality than v4 across this particular locus, and indeed our findings below illustrate that v4 contains genuine gene features that were lost during subsequent reassemblies.

### Protein analysis reveals sexual dimorphism in urinary MUP expression

Urinary MUPs are synthesized in the liver, secreted into the bloodstream and passed through the glomerular filter before being released in the urine (for a review [37]). Hepatic expression of MUPs is under sex and growth hormone control in both the mouse and the rat [38–40]. However, a striking difference between mouse and rat is the much more pronounced sexual dimorphism in expression of urinary MUPs in the rat. Whilst female mice have urinary MUP output that is approximately a third to a quarter that of males on average [8], female rats express virtually no MUPs in the liver [41–44] and as a consequence, no MUPs are apparent in urine [29, 30, 45].

The overall workflow for MUP characterization is summarized in Supplementary Figure 1. First, we measured the protein concentration in urine from male and female adults of two laboratory strains (Wistar Han and Brown Norway) as well as from some wild caught individuals. To correct for variation in urine dilution, protein output was normalized to urinary creatinine, a non-enzymatic by-product of muscle metabolism [46]. This confirmed that males had significantly higher protein output (Figure 2A). For Brown Norway males, the level was almost three times higher than that in females; for Wistar Han males it was almost five times higher and among wild males total urinary protein output was double that of females.

**Figure 2.**
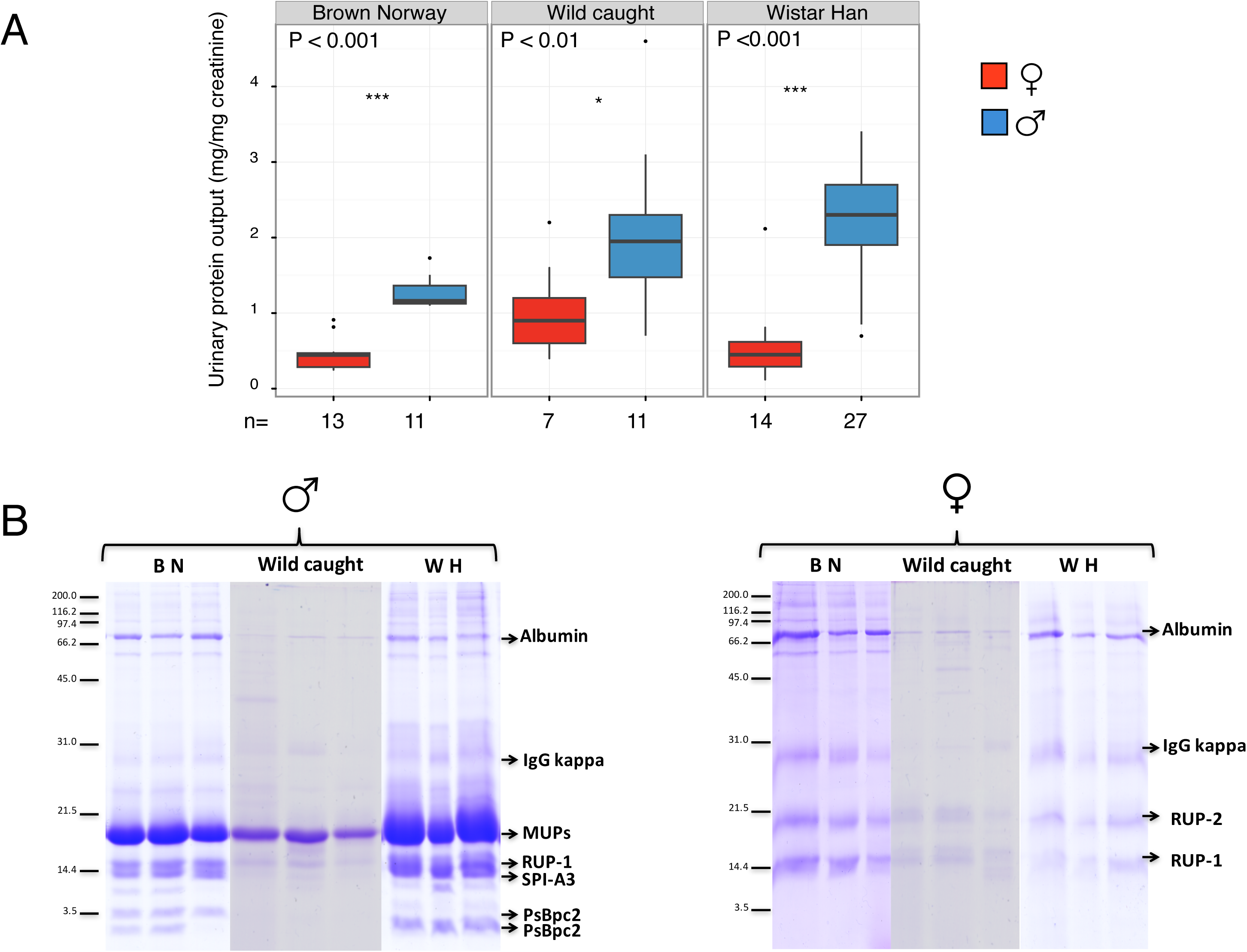
Protein expression in the urine of male and female rats. Urine samples were recovered from male and female rats of two different laboratory strains (Wistar Han, WH; Brown Norway, BN) and wild caught individuals. A: Protein output was expressed as mg protein/mg creatinine to correct for urine dilution. B: Urine samples were also analysed by SDS-PAGE. Proteins identified by in-gel digestion followed by PMF and tandem mass spectrometry are labelled and described in the text.

SDS-PAGE analysis of urine (Figure 2B) revealed a strongly expressed band at 18–19 kDa, present in all male samples and representing approximately 70 – 80 % of the total urinary protein. By contrast, a much fainter band was evident at a similar position in female urine. In-gel digestion, followed by PMF and tandem mass spectrometry, confirmed the presence of MUP peptides in the male expressed band, however, in-gel digestion revealed the identity of protein in the female band as rat urinary protein 2 (RUP-2, Uniprot KB P81828). The absence of peptides from MUPs in the female band is in good agreement with previous studies showing the scarcity of MUPs in female urine [29, 30, 45]. Additionally, in males, other proteins, including prostatic steroid binding protein (PsBpc2, Uniprot KB P02781) and the serine protease inhibitor A3K (SPI-A3, Uniprot KB P05545), were identified in other bands (Figure 2B). The urine of both sexes contained albumin, immunoglobulins and rat urinary protein 1 (RUP-1, Uniprot KB P81827, not to be confused with MUPs). Both laboratory rat strains and wild rats exhibited a low level of albuminuria.

### Phenotypic profiling to evaluate MUP heterogeneity and polymorphism

In laboratory mouse strains, the pattern of MUP expression is limited by inbreeding and consequent homozygosity at the *Mup* locus, but in wild-caught mice, heterogeneity is much more pronounced, both between mouse populations and between individuals of the same population [5, 6, 25, 47]. The highly polymorphic combinatorial nature of wild mouse urinary MUPs is the basis for individual recognition [10, 11, 13], driving assessment of genetic heterozygosity [48] and avoidance of inbreeding [12, 49]. It was of interest therefore to explore the heterogeneity in rat urinary MUPs. We have previously used ESI-MS to profile the isoforms of the MUPs secreted in mouse urine [6, 13, 25]. MUPs yield strong signals on ESI-MS and the intact masses can be determined to within ± 1 Da, permitting matching to predicted mature protein masses from genomic or cDNA sequences [25]. The masses obtained by ESI-MS correspond to the neutral average mass of the mature form of the protein, after the removal of the predicted signal peptide [50], and subtraction of 2 Da for the formation of a single disulphide bond, based on homology with known MUP structures [19, 36]. ESI-MS also allows semi-quantitative assessment of the relative amounts of each isoform [51].

In ESI-MS analysis of male urine, MUPs dominated the deconvoluted mass spectra (Figures 3 and 4) and several proteins and multiple discrete masses in the 18–19 kDa mass range were evident. All laboratory male rats, irrespective of strain, expressed proteins of masses 18712 Da, 18728 Da, and 18826 Da, the protein at 18728 being the most intense in all instances. Additionally, we identified strain-specific proteins at 18340 Da, 18420 Da and 18670 Da in Brown Norway rats, whereas proteins at 18553 and 18633 Da were exclusive to Wistar Han rats, the pattern being very stable within individuals of the same strain (Supplementary Figure 2).

**Figure 3.**
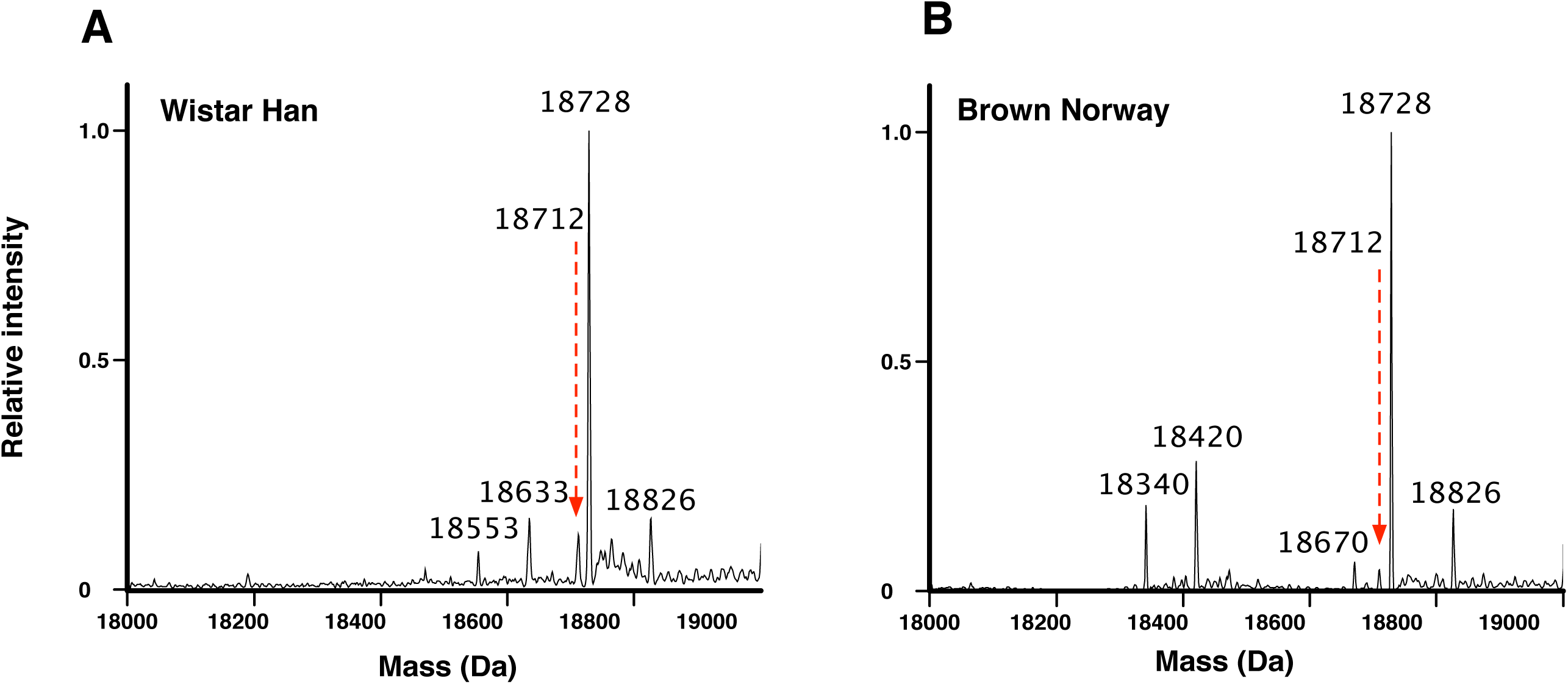
Intact mass protein profiling of male rat urine. Urine samples from Wistar Han (Panel A) or Brown Norway (Panel B) male rats were analysed by ESI-MS to obtain the profile of protein masses, here focused on 18,000 Da to 19,000 Da. Each spectrum is an average spectrum from 10 individual animal/urine replicates. Full data are presented in Supplementary Figure 2.

**Figure 4.**
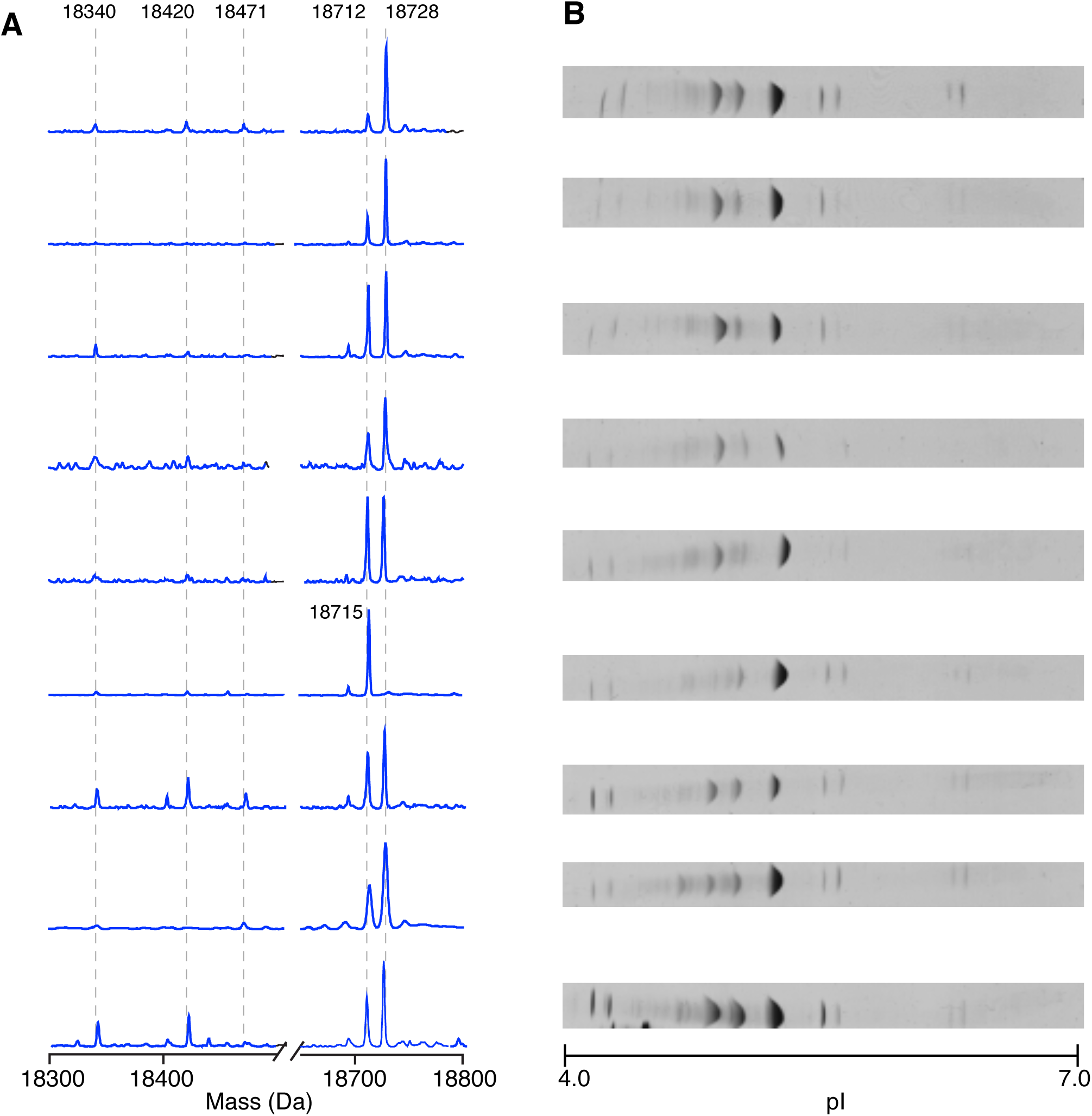
Intact mass protein profiling in wild male rat urine. Urine samples from male wild caught rats were analysed by ESI-MS intact protein mass profiling (Panel A) and by isoelectric focusing (Panel B).

In wild caught animals, urinary MUPs at 18728 Da and 18712 Da were dominant in 8 out of 9 individuals examined (Figure 4A). Only one individual was distinct in having a dominant peak at 18715 Da. Less abundant protein peaks were present, most prominently at 18340 Da and 18420 Da in six of the wild caught individuals and further minor peaks were observed at 18471 Da and 18694 Da. Thus, the pattern for wild individuals matched more closely that of the Brown Norway strain (Figure 3).

Although MUP profiles differed between the two laboratory strains and, as expected, within each strain the pattern was rather stable, the low degree of polymorphism among wild caught individuals (Figure 4A) was unanticipated. Compared with previous observations of house mice [5, 6, 13, 25, 47], there was significantly less polymorphism in protein isoforms as evidenced by the ESI-MS pattern of wild rats. To explore this in more detail, the same samples of wild rat urine were resolved by isoelectric focusing (IEF) (Figure 4B) to separate proteins by net charge – since MUPs were the predominant bands, they would be most prominent bands after isoelectric focusing. The protein banding patterns of nine individual male wild rats were similar and most of the urine samples resolved to three major and a few low intensity discrete bands. This was consistent with previous IEF studies of laboratory rat MUPs [22, 30, 52] but with fewer bands than recorded for house mice [10].

Roberts et al. [13] showed that mice are sensitive to changes in the relative ratios of MUP isoforms in urine. In rats, despite the absence of qualitative polymorphism between urine samples, the relative amount of each isoform differed between individuals. We quantified the relative abundance of each MUP mass from the peak area of the ESI-MS deconvoluted spectra, and calculated the correlation between the amounts of each protein, per individual (Figure 5). While laboratory strains showed high correlations between the relative amounts of the isoforms, this was not the case for samples derived from wild caught rats. The two protein masses that correlated in intensity most strongly among the wild caught individuals was the pair at 18340 Da and 18420 Da.

**Figure 5.**
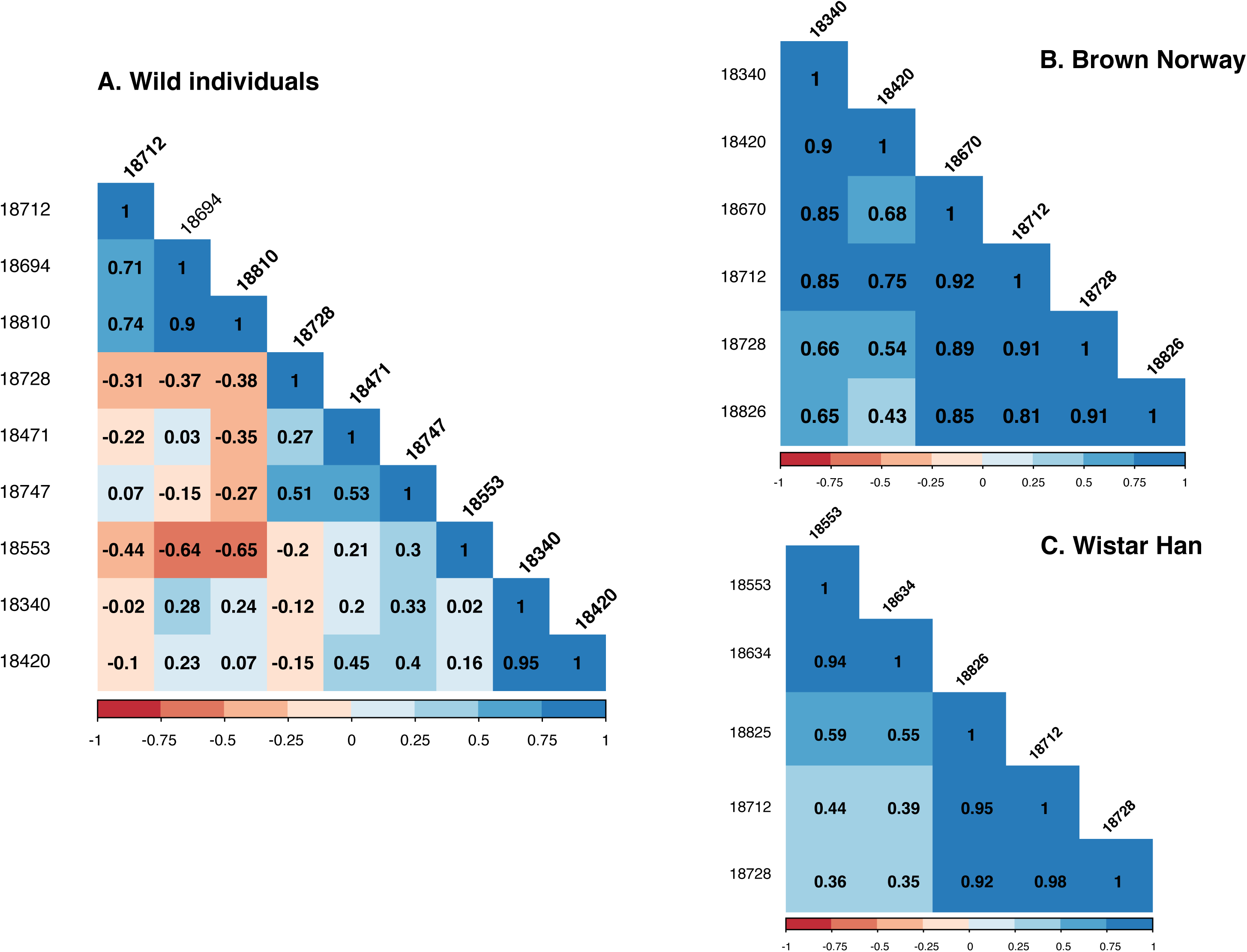
Spearman correlation coefficient analysis of intact mass areas. The relative amount of each isoform was quantified based on the peak area of the deconvoluted spectra. Spearman correlation coefficient was calculated between the amounts of each protein in different individuals. High correlation was found in laboratory strains whereas weak correlation was found in wild individuals, suggesting the possibility of quantitative polymorphism and a higher degree of variance than in the laboratory strains.

By contrast, ESI-MS of female rat urine (Supplementary Figure 3), showed two clusters of protein masses of around 11 kDa, and no mass peaks in the expected range of MUPs (18–19 kDa). We have not investigated these 11 kDa proteins further, but they are likely to be RUPs (rat urinary proteins). As anticipated, ESI-MS provides further confirmation of the lack of MUP expression in female rats.

### Characterization of the MUP proteoforms secreted in rat urine

To provide further MUP characterization, native gel electrophoresis and strong anion exchange fractionation were used to resolve the MUP mixture into discrete proteins to sequence by PMF and LC-MS/MS. For this purpose, we created an in-house database containing the sequences of the mature forms of MUPs derived from the gene annotation and transcript sequences published to date (Figure 1, Table 1), combined with all the protein entries in the Uniprot database for *Rattus norvegicus*. Supplementary Figures 4-8 provide the results of the different experimental approaches for the predicted proteins in Table 1. Supplementary Figure 9 shows a comparison of the protein sequences of the predicted rat MUP isoforms highlighting unique peptides for each isoform. From this detailed analysis, we could compile the evidence for each of the predicted proteins in the rat gene assembly.

**Table 1.**
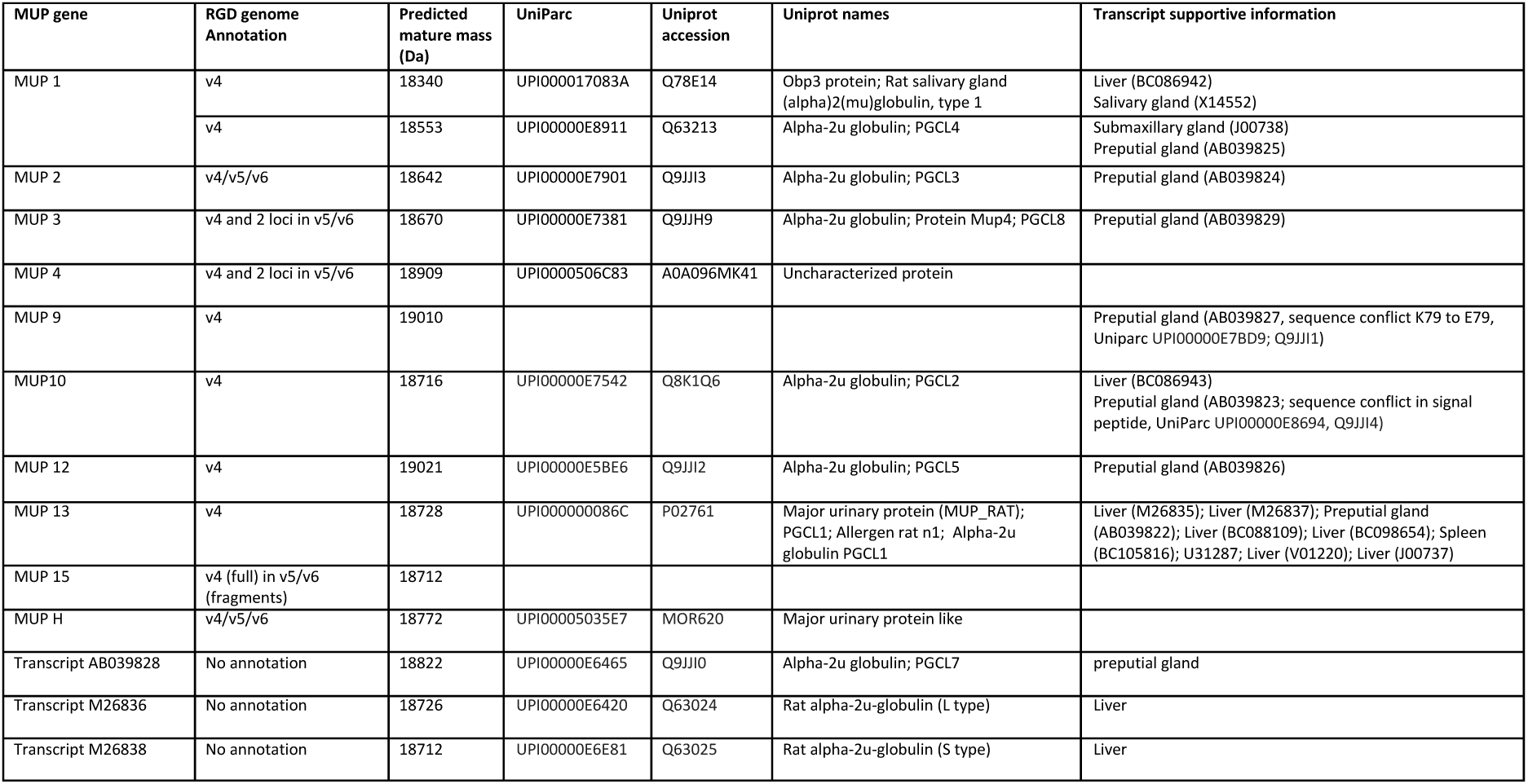
Current knowledge of rat MUP genes, transcript expression and protein products. This table shows an updated compilation of data from three releases of the rat genome sequence (RGSC_3.4 (v4) (December 2004), Rnor_5.0 (v5) (March 2012) and Rnor_6.0 (v6) (July 2014) from the Rat Genome Sequencing Consortium) using the annotations compiled in the Rat Genome Database (http://rgd.mcw.edu/) and Uniprot (http://www.uniprot.org/). Where possible, data relating predicted mature protein product is cross-correlated with experimental data that confirm true protein products.

In female rat samples, shotgun ‘bottom-up’ proteomics allowed the identification of MUPs at very low levels, in good agreement with previous papers establishing the presence of trace levels of MUPs in female urine [29, 30]. However, the very low abundance of these proteins meant that few peptides were observed and the resulting protein coverage did not allow confident assignment to any of the predicted proteins. By contrast, the same approach revealed the MUP isoform composition of male samples. Below, we discuss the protein-level evidence for each of the genes and include information from transcripts published to date (Table 1), focusing predominantly on genome assembly v4 for these assignments. For these analyses, we have retained the rat MUP numbering scheme first proposed by Logan et al [26] although this scheme also numbered the pseudogenes in the same sequence, in genome order. This numbering is now referenced in other studies [22, 53]. Indeed, a logical nomenclature based on gene order is impossible until a fully assembled and annotated analysis of this region of the rat genome is available.

#### Mup1 gene

Manual annotation of the genome assembly v4 predicts a protein of mature mass 18340 Da although, as previously mentioned, this gene was omitted from later assemblies (Figure 1). There are multiple transcriptional support data for this gene from liver (BC086942) [54], which is the primary source of urinary MUPs, and also salivary gland [55], with the cDNA predicting 18340 Da as mature protein mass. This MUP has also been referred to as OBP3 [22, 56], despite its high similarity to other MUPs and much lower similarity to rat OBP1 (28%) or OBP2 (18%). It is now clear that the gene encoding this protein is part of the MUP gene cluster. Further, unlike nasal MUPs in mice, which seem to be tissue specific, it is possible that the same MUP could play a dual role in odour reception and scent signalling, as it is expressed at high level in both the nose and urine. ESI-MS intact mass phenotyping showed the mass of 18340 Da in Brown Norway and in wild individuals. More detailed molecular analysis allowed assignment of this mass to the protein predicted by the *mup1* gene. Native gel electrophoresis followed by PMF provided good coverage of the protein (Supplementary Figure 5, band E and Supplementary Figure 6, band F). Besides, fractionation of the urine followed by proteolytic digestion of the protein and tandem MS of the peptides provided confident identification of the MUP1 protein (Supplementary Figure 8). There is also transcriptional support for the *mup1* gene from preputial gland (PGCL4) [57] and submaxillary gland (J00738; UniProt Q63213_RAT) [58]. However, after detailed examination of these sequences we conclude that they predict a protein identical to MUP1 except for two additional amino acids (-RG) at the C-terminus. This longer form predicts a mature mass of 18553 Da, a mass that we observed in the intact mass profile of Wistar Han males and occasionally in wild animals. Our analysis (Supplementary Figure 4, band E) allowed the assignment of the 18553 Da mass to the protein predicted by these transcripts, even though a gene designation may not have been possible because the Brown Norway strain used for the rat genome analysis does not express this mass.

We observed two protein peaks, of masses 18420 Da (Brown Norway and wild) and 18633 Da (Wistar Han) that could not be predicted from any of the genes described in any annotation of the MUP gene cluster, nor could these masses be generated by exopeptidase trimming of any known MUP sequence. Notably, these masses both differed from predicted masses 18340 Da and 18553 Da by 80 Da, a mass shift that might have been a consequence of multiple primary sequence changes but which was also consistent with the addition of a single phosphate group to a side chain residue. When urinary proteins were fractionated to resolve additional variants, the pairs at 18340/18420 Da, and 18553/18633 Da, eluted very closely in the chromatogram (Supplementary Figures 7 and 8), although the heavier protein was slightly more anionic in both cases, consistent with phosphorylation. Proteomic analysis of the chromatographic fractions containing proteins at 18633 Da and 18420 Da yielded extensive coverage for the protein sequences of that corresponded to the MUPs of masses 18553 Da and 18340 Da respectively, indicating a strong primary sequence relationship between the 80 Da separated proteins. To explore this further, LC-MS/MS peptide data from each protein fraction were analysed using Peaks software (Bioinformatics solutions Inc.) to search for post-translational modifications. For both protein fractions, the top-scoring endopeptidase Lys-C peptides revealed convincing evidence for phosphorylation of a serine residue at position 4 in the mature sequence of the protein (Figure 6). The proteins of masses 18553 Da and 18340 Da both share the same N-terminal sequence (Supplementary Figure 9). Manual annotation of the product ion spectra from either the Lys-C cleaved ([M+2H]^2+^, m/z=844.86) or tryptic N-terminal peptide ([M+2H]^2+^, m/z=474.18^2+^) revealed high quality coverage and unambiguous identification of a phosphorylation event at Ser_4_ (Figure 6). The phosphorylated forms were also resolved and identified from native gel electrophoresis and PMF (the unmodified N-terminal Lys-C peptide corresponding to m/z=1608 Da ([M+H]^+^) and the phosphorylated version corresponding to m/z=1688 Da ([M+H]^+^)) (Supplementary Figures 4, 5 and 6).

**Figure 6.**
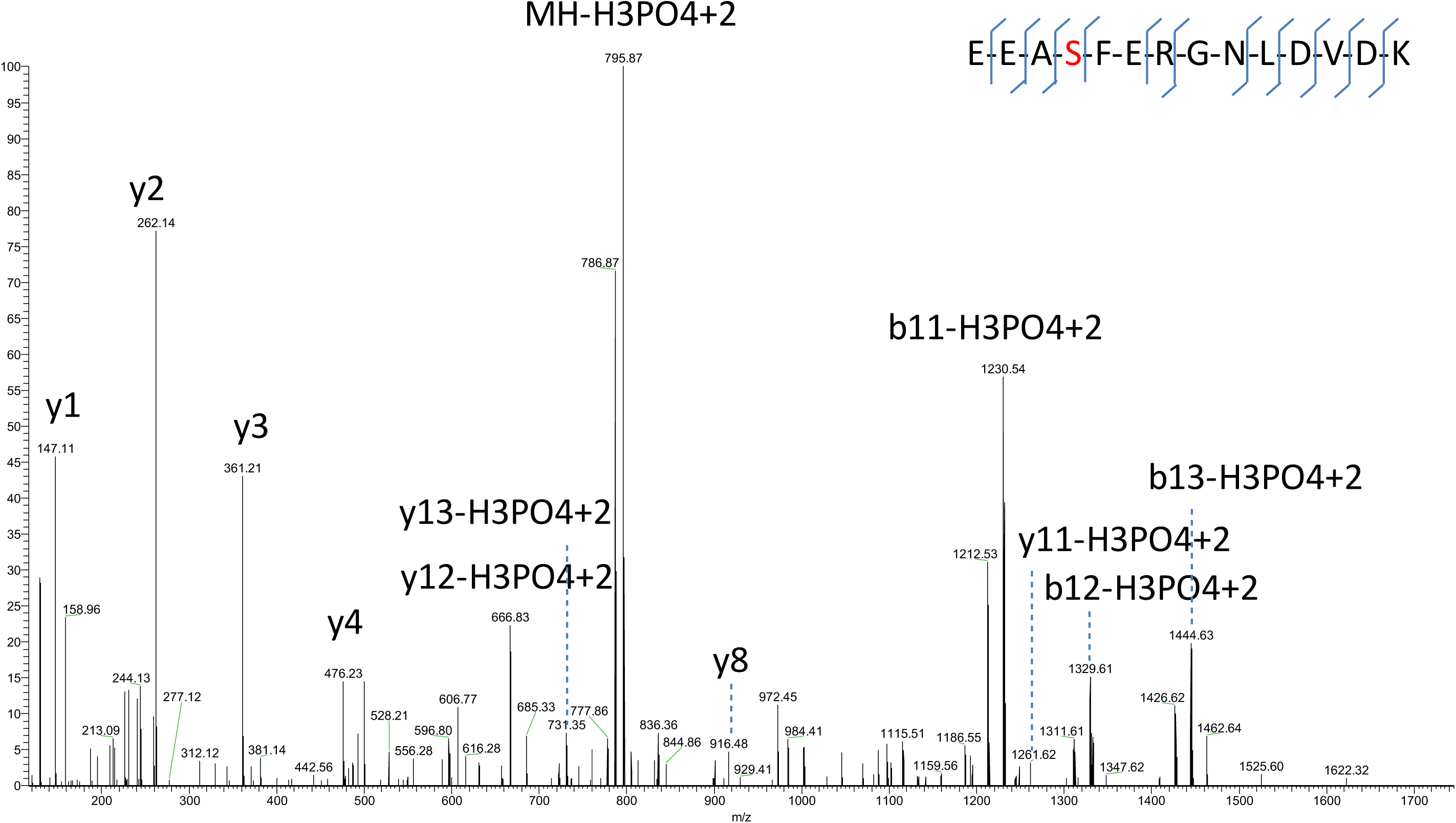
Evidence of phosphorylation of specific rat MUPs. Manual annotation of the N-terminal peptide of MUP1 showing evidence for phosphorylation of a serine residue at position 4.

Although phosphorylation of extracellular proteins is not well studied, the Ser_4_ residue sits within a consensus sequence motif (SxE) (Supplementary Figure 9) for phosphorylation by FAM20C kinase, the enzyme responsible for the phosphorylation of most secreted proteins in humans [59]. The rat genome contains an ortholog gene of human FAM20C kinase on chromosome 12 (RGD:1311980) and other members of the rat MUP family contain this phosphorylation motif (Supplementary Figure 9), invoking the possibility of phosphorylation in other isoforms, although we have no evidence so far that other isoforms are phosphorylated to the same extent as MUP1.

There is a report of phosphorylation of MUPs in *Rattus rattus*, specifically from the preputial gland [60]. Phosphorylation of MUPs was considered based on spot distribution on 2D gels, however no other evidence was provided and the proposed site, at Ser51 does not sit within the consensus sequence of FAM20C kinase. Therefore, this putative phosphorylation site requires further validation. Phosphorylation significantly influences ligand binding affinities of porcine OBP [61] suggesting that this may have an influence on both the signature of urinary volatiles bound and released by MUPs and the capture of odours in the nose, but further studies are needed to understand the significance of this modification.

#### Mup2 gene

The gene encoding this protein predicts a mature mass of 18642 Da. There is transcriptional support for this protein sequence from rat preputial gland (PGCL3, [43]). We found no evidence for this mass in intact mass profiles of either intact urine or after ion exchange fractionation. However, shotgun proteomics gave us high protein coverage including peptides unique to MUP2: 80% protein coverage in both Wistar Han and wild individuals; and 60% protein coverage in Brown Norway. Therefore, we hypothesized three possibilities for the absence of the 18642 Da mass in the intact mass profile: the protein is only present in small amounts, the protein is phosphorylated, or the protein is trimmed at the N-terminal. Regarding phosphorylation, although this protein contains a serine residue at mature sequence position 4, it does not contain the sequence motif for FAM20C kinase and, as anticipated, there was no evidence for phosphorylation in the N-terminal peptide in shotgun proteomics analysis. A common feature in all the identifications of this protein, regardless of individual donor, was the incomplete coverage of the N-terminal part of the sequence, which might suggest trimming of the N terminus to remove between 7 and 10 amino acids. However, we were unable to identify an intact mass peak matching a trimming event either.

#### Mup3 gene

The predicted protein mass for *mup3* gene product is 18670 Da. Again, there is transcriptional support from rat preputial gland (PGCL8 [43]). PMF after native electrophoresis allowed the identification of this protein in Wistar Han (Supplementary Figure 4, band B), but not in Brown Norway or wild individuals. Further, shotgun proteomics provided 66% sequence coverage and 5 unique peptides on average. However, no evidence of this mass was obtained in the intact mass profiling, suggesting that the protein is expressed at low levels only. The intact mass profile of Brown Norway and wild individuals showed a peak at 18670 Da, however, protein fractionation and further analysis demonstrated that this mass does not correspond to MUP3 but likely to a trimming of the 18728 Da form at the C-terminal (-G) (discussed below).

#### Mup4 gene

The predicted protein mass for *mup4 gene* product is 18909 Da. There is no transcriptional support for this gene, and we could find no evidence for a gene product in urine in any of the individuals. However, some MUPs are not expressed in urine, and we cannot exclude the possibility of expression in a tissue other than liver, the likely source of urinary MUPs.

#### Mup9 gene

For this sequence, SignalP [62] predicts a signal peptide two amino acids shorter than that commonly observed in MUPs (17 instead 19 amino acids). Hence this isoform is two amino acids longer than the rest of the isoforms at the N-terminus (Supplementary Figure 9) and the predicted mass of the mature protein is 19010 Da. No peak at that mass was found in any of the samples that were analysed. By contrast, Wistar Han and some wild individuals demonstrated a mass peak at 18745 Da, which matches the predicted protein mass of the *mup9* mature gene product after removal of the usual 19 amino acid signal peptide. Furthermore, while no evidence was found for the predicted N-terminal peptide corresponding to the long form (HEEEASFER-), the N-terminal peptide corresponding to the ‘short form’ (EEASFER-) was readily identified by PMF and LC-MS/MS after native PAGE and in-gel digestion of the corresponding band (Supplementary Figure 4, band A). Therefore, we venture that in this instance, the prediction of the signal peptide cleavage is incorrect and that in common with other MUPs, this protein loses a signal peptide of 19 amino acids and has the commonly seen N-terminal sequence of GluGlu. Further, a minor sequence conflict arose at Lys_81_ in the annotated sequence of the *mup9* gene from genome assembly v4, to Glu_81_ suggested by the transcript AB039827 (PGCL6 [43]). We confirmed by in-gel digestion after native PAGE that Wistar Han males possess the Glu_81_ form. Additional evidence is provided by shotgun proteomics, yielding good coverage for this protein in Wistar Han and some wild individuals and equally confirming the Glu_81_ form. There was no evidence for the expression of this protein in the Brown Norway strain.

#### Mup10 gene

The predicted protein mass for *mup10 gene* product is 18716 Da, supported by transcriptional information from rat liver and preputial gland (PGCL2 [43]). Intact mass analysis, in-gel digestion after native PAGE and SAX fractionation approaches all provided conclusive evidence for the *mup10* predicted protein sequence in both laboratory strains and wild individuals (Supplementary Figure 4).

#### Mup12 gene

The predicted protein mass for *mup7 gene* product is 19021 Da, for which there is transcriptional support from rat preputial gland (PGCL5 [43]). However, we did not find confident evidence for the expression of this protein in any of the urine samples analysed.

#### Mup13 gene

The predicted protein mass for *mup13 gene* product is 18728 Da with multiple transcripts supporting the expression of this gene (Table 1). For both laboratory strains, and for wild caught individuals, we obtained confident identification of the protein predicted by the *mup13* gene. This mass is the most intense in the ESI-MS profile (Figures 3 and 4) and the most intense in native or IEF electrophoresis (Supplementary Figure 4 and Figure 4).

Some of the observed masses in the ESI-MS profile are consistent with an N-terminally processed protein of 18728 Da. For example, the mass at 18470 Da, observed in the ESI-MS protein profile in some individuals, is consistent with trimming of the N-terminal amino acids from the 18728 Da isoform. SAX fractionation revealed three masses in the flow through volume (18470, 18399 and 18312 Da) that can be explained by the trimming of the N-terminal amino acids from the 18728 Da form (EE-, EEA- and EEAS-, respectively). The trimming of these amino acids means that the pI becomes close to 6 for all three proteins, which is the pH at which the chromatography is performed and explains their appearance in the flow through (the net charge of the proteins is zero under these conditions, preventing the protein from binding to the column). Native electrophoresis allowed the isolation and sequencing of the protein corresponding to the predicted mass 18470 Da, confirming the trimming of the N-terminus (Supplementary Figure 4-6, band A). For this protein, we identified the N-terminal Lys-C cleaved peptide, corresponding to the removal of EE-, at 1217.58 Da by both PMF and LC-MS/MS. Another example of trimming is the protein explaining the mass 18670 Da, seen specifically in the ESI-MS from Brown Norway rats (Supplementary Figure 5, band B). We also found evidence for a C-terminal trimming of the 18728 Da protein (loss of a glycine residue) that would explain the mass 18670 Da (within 1 Da instrument error). Although the MUPs in rodent urine are generally proteolytically-resistant, rat urine contains proteases that could attack the termini of the protein (such as meprin and neprilysin [63], Gómez-Baena et al, in preparation), although further experiments are needed to explore the extent of processing in urine and the biological significance thereof.

In one wild-caught individual, the mass at 18728 Da was less intense in the ESI-MS profile (Figure 4A, individual 6L), although the band in the range of the 18728 Da in the IEF profile (Figure 4B) was strongly stained for this sample. PMF of this IEF band allowed the sequencing of a new MUP sharing the sequence of the 18728 form but with one amino acid change, from Thr to Ser in position 154 which corresponds with a mass shift of -14 Da explaining the mass of 18714 Da in the intact mass profile for this individual. This mutation was also confirmed by MS/MS data using Peaks PTM predictor.

#### Mup15 gene

The predicted protein mass for *mup15 gene* product is 18712 Da. There is no transcriptional support for expression and no protein of this mass was apparent in rat urine. Although the intact mass profile shows a peak at 18712 Da, further analysis showed that this mass does not correspond to the predicted MUP15 protein sequence, but the sequence of the transcript M26838 (discussed below).

#### MupH gene

We refer to this as *mupH* to reflect the fact that it was only identified in later genome assemblies and was thus labelled. The predicted protein mass for *mupH gene* product is 18772 Da. There is no transcriptional support for expression of this gene and there was no evidence for the expression of this protein in any of the samples analysed. It is not yet clear whether this is a true protein coding gene.

AB039828 transcript: This transcript was isolated from preputial gland (PGCL7 [43]) and would have a predicted mass for the mature protein of 18822 Da. Although we were not able to identify the mass 18822 Da in the ESI-MS profile, the mass of 18694 Da, seen in some wild males, matches the cleavage of a single E from the N-terminal of the 18822 Da MUP (18693.4 Da). However, we could obtain no data to support this possibility.

M26836 transcript: This transcript was isolated from liver [54]. The predicted mass for the protein is 18726 Da. However, we could find no evidence for expression of this protein in urine.

M26838 transcript: This transcript was also isolated from liver [54]. The predicted mass for the protein is 18712 Da. In the ESI-MS profile of most of the males a mass at 18712 Da was observable. Native gel electrophoresis followed by PMF and LC-MS/MS allowed confident identification of the protein predicted by the M26838 transcript in wild individuals (Supplementary Figure 4). This protein is one of the most intense bands in the native gels and is likely to be a highly expressed MUP in wild individuals, while our results suggest a lesser expression in Wistar Han and Brown Norway strains.

### Analysis of the protein products of the gene cluster

We provide a detailed analysis of the isoforms of the major urinary protein system expressed in the urine of *Rattus norvegicus*. We characterized the urinary MUPs from two of the most widely used laboratory strains, Wistar Han and Brown Norway, as well as wild caught individuals. We provide evidence at the protein level for several proteins predicted by the genome assembly suggesting strain-specific expression. There are two levels of variance: at the gene and allele level and at the post-translational level. The entire panoply of the urinary protein products, and their post-translational space, is summarized in Figure 7. Most of the residues that differ between MUP10, MUP13 and M26838 are not shared with other rat MUPs (E_4_D in MUP10, D_15_A in MUP13, L_118_A in MUP10 and M26838, R_158_H in M26838), although one variant (D_29_N) is shared across MUPs 10, 13, 4 and H. The degree of similarity between these three rat MUPs is similar to that between a set of highly similar MUPs in house mice encoded by approximately 15 genes in the central region of the mouse MUP cluster [25]. In mice, these highly similar MUPs provide the basis for an individuality signal in urine scent marks [10, 13, 14], with each individual expressing a fixed signature of these MUPs. Combinatorial polymorphism arises both from variation in MUP sequences (involving a limited set of variable sites) and differential transcription of Mup genes [25]. Mice are able to discriminate different MUP signatures, both through V2Rs in the vomeronasal organ that detect MUPs directly [14] and through differences in the signature of ligands bound and released by MUPs [13]. Although rats express fewer MUPs in urine than mice, combinatorial polymorphism in the relative amounts of each MUP could still provide considerable capacity to encode individual differences. Consistent differences in MUP signatures between strains suggest a high degree of genetic determination, but studies have not yet addressed how stable MUP profiles are in rats, or the sensitivity of rats to discriminate these relatively small differences between rat MUP isoforms or their relative ratios. The molecular characterization of the MUP proteoforms expressed by rats presented here now provides the opportunity for such detailed studies to be carried out, an essential next step to understand the functions of MUPs in rat scent signals. Understanding whether some MUPs, or the extent of post-translational modifications, are particularly sensitive to the hormonal and/or behavioural status of individuals could also provide very useful insight into potential functions.

**Figure 7.**
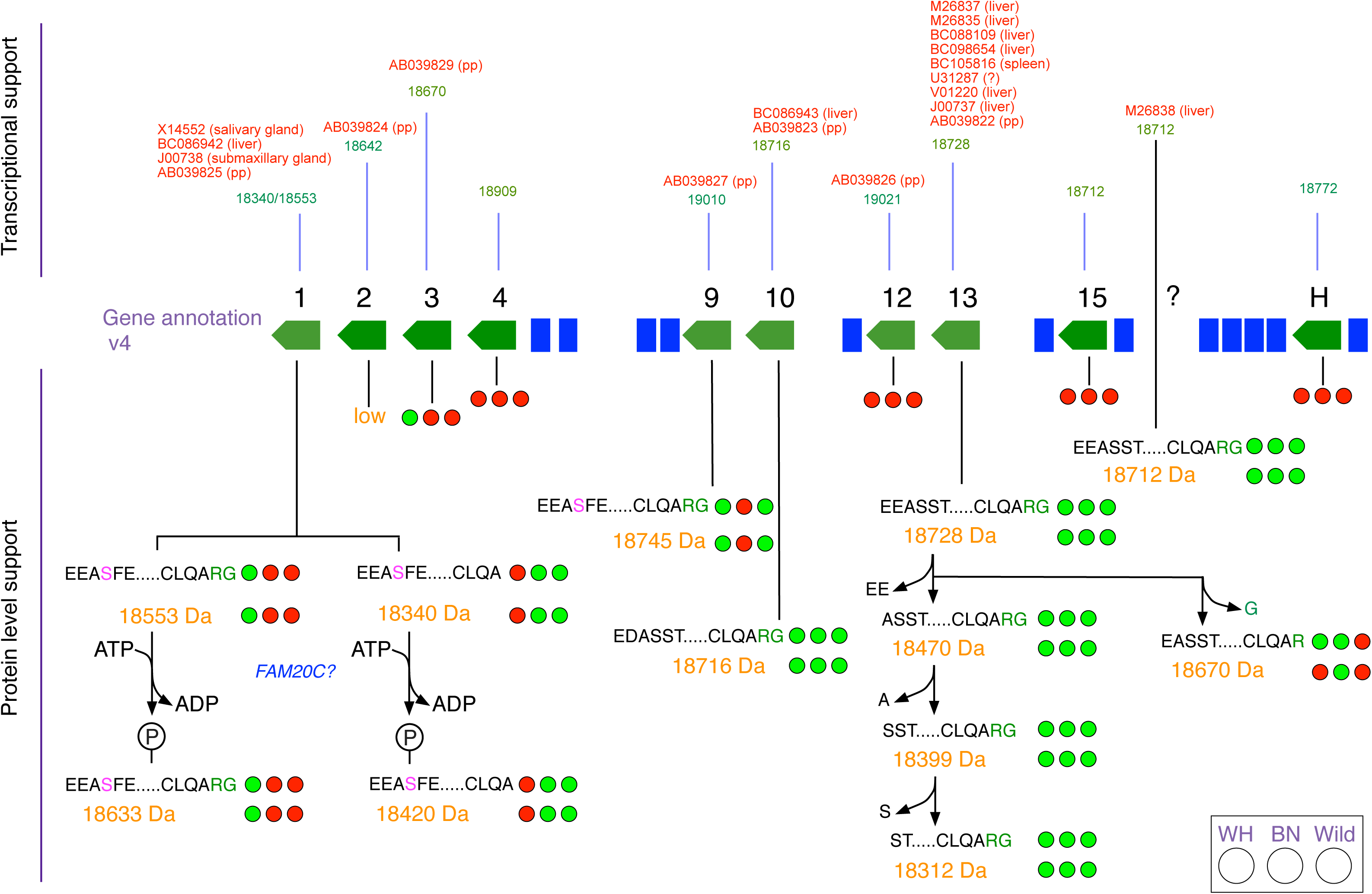
Summary of phenotypic profiling of rat urinary MUPs. The phenotypic analysis of urinary MUPs is mapped to the *mup* region of rat genome version 4. Above the gene annotation we include transcriptional evidence from other studies, highlighting tissue of origin. Below the gene annotation, we summarise the findings in this paper. For each protein, we report the evidence for a mature gene product by intact mass analysis (adjacent to the predicted mass in orange) and from bottom up peptide analysis (adjacent to the sequence data) for each of the three groups of animals tested: Wistar Han (WH), Brown Norway (BN) and wild-caught (Wild) individuals. A green circle defines confident protein-level evidence, a red circle denotes the absence of evidence for this particular gene product.

Most strikingly, our analysis further revealed the complexity of the post-translational modifications that are applied to rat MUPs, including phosphorylation of MUP1 and protein trimming of MUP8 (summarized in Figure 7). Neither of these modifications is evident in the best studied MUP system in the mouse, *Mus musculus domesticu*s and it is possible that the rat relies on post-translational modification to elicit further variance in semiochemical properties, but confirmation of this must await functional bioassay in behavioural tests. Additionally, our study emphasizes the need for detailed protein analysis to identify individual proteoforms prior to functional characterization.

## MATERIALS AND METHODS

### Animals and urine collection

Laboratory rat urine donors were 11 Wistar Han^®^ outbred rats and 11 Brown Norway BN/RijHsd inbred rats obtained from Harlan UK (now Envigo) at 3 weeks of age. Animals were housed in GPR2 cages (56 x 38 x 25 cm, North Kent Plastics, UK) on Corn Cob Absorb 10/14 substrate (IPS Product Supplies Ltd, London). Water and food (lab diet 5002, Purina Mills) were given *ad libitum.* All rats were provided with paper wool nesting material, cardboard houses and plastic tubes (8 cm diameter) for home cage enrichment. Urine samples were obtained from adult rats aged 3 to 9 months. For urine collection from laboratory strains, individual rats were placed in a clean empty wire-floored polypropylene RC2R cage (56×38×22cm) without food or water. The cages were suspended over trays (checked every 30 min) into which the urine could collect. After 2–4 h rats were returned to their home cage. Adult wild rat samples were provided by the former Central Science Laboratory of Defra (now part of the Animal and Plant Health Agency, UK) from rats that were trapped on farms within 15 miles of the Central Science Laboratory (Sand Hutton, North Yorkshire). Wild-caught animals were individually housed in suspended wire cages with free access to food and water. Urine samples were collected overnight on a clean waxed paper sheet in the tray under the cage. All samples were aspirated by pipette, avoiding feces and food fragments, and stored at -20 °C until use.

### Protein and creatinine concentration assays

Protein concentration was determined using the Coomassie Protein Plus assay kit (Thermo Scientific). Urinary creatinine was quantified using a creatinine assay kit (Sigma-Aldrich).

### Electrospray ionization mass spectrometry (ESI-MS) of intact proteins

Urine samples were diluted in 0.1% (v/v) formic acid and centrifuged at 13,000 g for 10 min. All analyses were performed on a Synapt G2 mass spectrometer (Waters Corporation), fitted with an API source. Samples were desalted and concentrated on a C4 reverse phase trap (Thermo Scientific) and protein was eluted at a flow rate of 10 μL/min using three repeated 0–100 % (v/v) acetronitile (ACN) gradients. Data was collected between 800 and 3500 Th (m/z), processed and transformed to a neutral average mass using MaxEnt 1 (Maximum Entropy Software, Waters Corporation). The instrument was calibrated using a 2 pmol injection of myoglobin from equine heart (Sigma-Aldrich; M1882).

### Polyacrylamide gel electrophoresis (PAGE)

SDS-PAGE was performed as described by Laemmli [64]. Samples were resuspended in 2x SDS sample buffer (125 mM Tris-HCl; 140 mM SDS; 20% (v/v) glycerol; 200 mM DTT and 30 mM bromophenol blue) and heated at 95 °C for 5 min before loading onto gel. Electrophoresis was set at a 200 V constant potential for 45 min through a 4 % (w/v) stacking gel followed by a 15 % (w/v) resolving polyacrylamide gel. PAGE under native conditions was performed following the same protocol but in the absence of SDS and DTT during the process. Electrophoresis was set at a 200 V constant potential for 60 min. Protein bands were visualized with Coomassie Brilliant Blue stain (Sigma-Aldrich).

### Isoelectric focusing (IEF)

IEF was performed using a Multiphor flatbed system (Amersham Biosciences) using an Immobiline dry-plate gel, pH range 4–7 (GE Healhcare Life Sciences) and cooled to 10 °C. Urine samples were concentrated and desalted using Vivaspin centrifugal concentrators (3 kDa MWCO, Vivascience). Urine samples were diluted to 1 mg/mL with deionized water and 5 μL was applied to sample strips placed on the gel. Samples were loaded into the gel at 200 V, 5 mA and 15 W for 200 V⋅ h. The sample strips were removed and the gel was run at 3500 V, 5 mA and 15 W for 14.8 kV⋅ h. After fixation with 20 % TCA (v/v), the gel was stained with Coomassie Brilliant Blue.

### Strong anion exchange chromatography (SAX)

Urine was desalted using Zeba columns (Pierce, 0.5 mL) and then filtered through a 0.45 μm Millipore filter prior to injection. Proteins in rat urine were separated in different fractions by high resolution strong anion exchange on an AKTA instrument equipped with a Resource Q column (GE Life Sciences, V= 1 mL). The column was equilibrated with MES buffer (50 mM, pH 6), and bound proteins were eluted using a linear gradient of 0 to 1 M NaCl over 20 column volumes, with a flow rate of 2 mL/min. Fractions were manually collected and analysed individually.

### In gel digestion

Gel plugs were removed from the gel using a Pasteur glass pipette, placed into low binding tubes and then destained using 50 μL of 50 mM ammonium bicarbonate/50 % (v/v) ACN for 30 min at 37 °C. The plugs were then incubated with 10 mM dithiothreitol (DTT) for 60 min at 60 °C. The DTT was then discarded and 55 mM iodoacetamide (IAM) stock solution was added to each tube and incubated for 45 min at room temperature in the dark. After discarding the IAM, the plugs were washed twice using 50 mM ammonium bicarbonate/50 % (v/v) ACN. The plugs were then dehydrated by adding 10 μL of 100 % ACN. Sequencing grade endoproteinase Lys C (Wako) (diluted in 25 mM Tris-HCl, 1 mM EDTA, pH 8.5) was then added and the digests incubated overnight at 37 °C. The reaction was stopped by adding formic acid solution to a 1 % final concentration (v/v).

### Peptide mass fingerprinting (PMF)

Peptide mixtures from the proteolytic digestion reactions were analysed on a Bruker UltraFlex matrix-assisted laser-desorption ionization–time of flight-mass spectrometer (MALDI– TOF) (Bruker Daltonics), operated in the reflectron mode with positive ion detection, or a MALDI Synapt G2 Si (Waters Corporation). Samples were mixed 1:1 (v/v) with a 10 mg/mL solution of α-cyano-4-hydroxycinnamic acid in 60 % ACN/0.2 % TFA (v/v), before being spotted onto the MALDI target and air-dried. Spectra were acquired at 35–40 % laser energy with 500-2000 laser shots per spectrum. Spectra were gathered between m/z 900 and 4500. External mass calibration was performed using a mixture of des-Arg bradykinin (904.47 Da), neurotensin (1672.92 Da), ACTH (corticotrophin, 2465.2 Da) and oxidized insulin â chain (3495.9 Da) (2.4, 2.4, 2.6 and 30 pmol/µL, respectively) in 50 % ACN/0.1 % TFA (v/v).

### In solution digestion

Liquid samples were denatured with RapiGest (Waters Corporation) and alkylated, prior to digestion with trypsin or endopeptidase Lys C. To stop the proteolytic reaction and to inactivate and precipitate the detergent, TFA (final concentration 0.5 % (v/v)) was added, followed by incubation for 45 min at 37 °C. To remove all insoluble material, samples were centrifuged twice at 13,000 g for 15 min [65].

### Liquid chromatography-tandem mass spectrometry analysis

LC-MS/MS analysis was performed using a QExactive instrument (Thermo Scientific) coupled to an Ultimate 3000 LC nano system (Thermo Scientific). Protein digests were resolved on an Easy-spray PepMap RSLC C18 column over a linear gradient from 3 to 40% (v/v) ACN in 0.1% v/v formic acid. The QExactive instrument was operated in data dependent acquisition mode. Full scan MS spectra (m/z 300-2000) were acquired at 70,000 resolution and the ten most intense multiply charged ions (charge ≥ 2) were sequentially isolated and fragmented by high energy collisional dissociation (HCD) at 30% standardized collision energy. Fragments ions were detected at 35,000 resolution and dynamic exclusion was set at 20 s. Proteome Discoverer (Thermo Scientific) version 1.4 was used to generate peak lists using default parameters and Mascot version 2.4 (Matrix Science) to identify peptides and proteins, using a database containing all the entries annotated for *Rattus norvegicus* in Uniprot (www.uniprot.org) (updated on 20170605) and the sequences of the mature MUP proteins, applying a FDR < 1 %. Either trypsin or Lys C was selected as the specific enzyme, allowing one missed cleavage. MS/MS data were also analysed using Peaks Studio 8.0 (Bioinformatics solutions Inc.) to identify post-translational modifications. All raw mass spectrometry files will be made immediately available upon request.

### Data analysis

Data were visualised and analysed using Aabel (Gigawiz software, http://www.gigawiz.com/) and R (v.3.2) (http://www.R-project.org/). Protein maps were generated using PeptideMapper [66].

## Supporting information

**Supplementary Figure 1** Workflow followed to analyze rat urine samples.

**Supplementary Figure 2** ESI-MS intact mass deconvoluted spectra from individual Wistar Han and Brown Norway males.

**Supplementary Figure 3** ESI-MS analysis of female rat urine. Deconvolution of the spectrum estimates two masses of about 11 kDa (11065 and 11450 Da) likely corresponding to the rat urinary proteins 1 and 2.

**Supplementary Figure 4** Sequencing of MUP isoforms from Wistar Han males by native electrophoresis of urine followed by in-gel LysC digestion and analysis by PMF and LC-MS/MS. Peptide maps show sequence coverage. Red boxes show unique peptides for the isoform and blue boxes show common peptides to several MUP isoforms.

**Supplementary Figure 5** Sequencing of MUP isoforms from Brown Norway males by native electrophoresis of urine followed by in-gel LysC digestion and analysis by PMF and LC-MS/MS. Peptide maps show sequence coverage. Red boxes show unique peptides for the isoform and blue boxes show common peptides to several MUP isoforms.

**Supplementary Figure 6** Sequencing of MUP isoforms from wild caught males by native electrophoresis of urine followed by in-gel LysC digestion and analysis by PMF and LC-MS/MS. Peptide maps show sequence coverage. Red boxes show unique peptides for the isoform and blue boxes show common peptides to several MUP isoforms.

**Supplementary Figure 7** Sequencing of MUP isoforms from Wistar Han males by ion exchange chromatography fractionation followed by LysC digestion and analysis by LC-MS/MS. ESI-MS intact mass deconvoluted spectra from individual Wistar Han are shown for each fraction.

**Supplementary Figure 8** Sequencing of MUP isoforms from Brown Norway males by ion exchange chromatography fractionation followed by LysC digestion and analysis by LC-MS/MS. ESI-MS intact mass deconvoluted spectra from individual Brown Norway are shown for each fraction.

**Supplementary Figure 9** Network representation of a comparison of the expected trypsin peptides from protein sequences of the predicted rat MUP isoforms, highlighting unique peptides for each isoform. Peptide mapper [66] was used to perform in-silico digestion of protein sequences and network was built using Cytoscape [68].

## Acknowledgements

This work is supported by the Biotechnology and Biological Sciences Research (BBSRC) programme BB/J002631/1. We are grateful to Dr. A D MacNicoll for supplying the wild rat samples. The authors gratefully acknowledge instrumental support from Dr. Philip Brownbridge (Centre of Proteome Research), Dr. Richard Humphries, Amanda J. Davidson, John Waters, Rachel Spencer and Joshua Beeston (Mammalian Behaviour and Evolution Group).

## References

1. Finlayson JS, Potter M, Runner CR. Electrophoretic Variation and Sex Dimorphism of the Major Urinary Protein Complex in Inbred Mice: A New Genetic Marker. Journal of the National Cancer Institute. 1963;31:91–107. Epub 1963/07/01. PubMed PMID: 14043041.

2. Roy AK, Neuhaus OW, Gardner E. Studies on Rat Urinary Proteins. Federation proceedings. 1965;24(2p1):507-&. PubMed PMID: ISI:A19656257802090.

3. Finlayson JS, Morris HP. Molecular Size of Rat Urinary Protein. Proceedings of the Society for Experimental Biology and Medicine Society for Experimental Biology and Medicine. 1965;119:663–6. Epub 1965/07/01. PubMed PMID: 14328970.

4. Robertson DH, Cox KA, Gaskell SJ, Evershed RP, Beynon RJ. Molecular heterogeneity in the Major Urinary Proteins of the house mouse Mus musculus. Biochem J. 1996;316 (Pt 1):265–72. Epub 1996/05/15. PubMed PMID: 8645216; PubMed Central PMCID: PMC1217333.

5. Robertson DH, Hurst JL, Bolgar MS, Gaskell SJ, Beynon RJ. Molecular heterogeneity of urinary proteins in wild house mouse populations. Rapid communications in mass spectrometry : RCM. 1997;11(7):786–90. Epub 1997/01/01. doi: 10.1002/(SICI)1097-0231(19970422)11:7<786::AID-RCM876>3.0.CO;2-8. PubMed PMID: 9161047.

6. Hurst JL, Beynon RJ, Armstrong SD, Davidson AJ, Roberts SA, Gómez-Baena G, et al. Molecular heterogeneity in major urinary proteins of Mus musculus subspecies: potential candidates involved in speciation. Scientific reports. 2017;7:44992. doi: 10.1038/srep44992. PubMed PMID: 28337988; PubMed Central PMCID: PMCPMC5364487.

7. Hurst JL, Robertson DHL, Tolladay U, Beynon RJ. Proteins in urine scent marks of male house mice extend the longevity of olfactory signals. Animal behaviour. 1998;55(5):1289–97. Epub 1998/12/16. PubMed PMID: 9632512.

8. Beynon RJ, Hurst JL. Urinary proteins and the modulation of chemical scents in mice and rats. Peptides. 2004;25(9):1553–63. Epub 2004/09/18. doi: 10.1016/j.peptides.2003.12.025. PubMed PMID: 15374657.

9. Kwak J, Strasser E, Luzynski K, Thoss M, Penn DJ. Are MUPs a Toxic Waste Disposal System? PLoS One. 2016;11(3):e0151474. doi: 10.1371/journal.pone.0151474. PubMed PMID: 26966901; PubMed Central PMCID: PMCPMC4788440.

10. Hurst JL, Payne CE, Nevison CM, Marie AD, Humphries RE, Robertson DH, et al. Individual recognition in mice mediated by major urinary proteins. Nature. 2001;414(6864):631–4. Epub 2001/12/12. doi: 10.1038/414631a. PubMed PMID: 11740558.

11. Cheetham SA, Thom MD, Jury F, Ollier WE, Beynon RJ, Hurst JL. The genetic basis of individual-recognition signals in the mouse. Current biology : CB. 2007;17(20):1771–7. Epub 2007/10/24. doi: 10.1016/j.cub.2007.10.007. PubMed PMID: 17949982.

12. Green JP, Holmes AM, Davidson AJ, Paterson S, Stockley P, Beynon RJ, et al. The Genetic Basis of Kin Recognition in a Cooperatively Breeding Mammal. Curr Biol. 2015;25(20):2631–41. doi: 10.1016/j.cub.2015.08.045. PubMed PMID: 26412134.

13. Roberts SA, Prescott MC, Davidson AJ, McLean L, Beynon RJ, Hurst JL. Individual odour signatures that mice learn are shaped by involatile major urinary proteins (MUPs). BMC biology. 2018;16(1):48. doi: 10.1186/s12915-018-0512-9-. PubMed PMID: 29703213; PubMed Central PMCID: PMCPMC5921788.

14. Kaur AW, Ackels T, Kuo TH, Cichy A, Dey S, Hays C, et al. Murine pheromone proteins constitute a context-dependent combinatorial code governing multiple social behaviors. Cell. 2014;157(3):676–88. doi: 10.1016/j.cell.2014.02.025. PubMed PMID: 24766811.

15. Chamero P, Marton TF, Logan DW, Flanagan K, Cruz JR, Saghatelian A, et al. Identification of protein pheromones that promote aggressive behaviour. Nature. 2007;450(7171):899–902. Epub 2007/12/08. doi: 10.1038/nature05997. PubMed PMID: 18064011.

16. Roberts SA, Simpson DM, Armstrong SD, Davidson AJ, Robertson DH, McLean L, et al. Darcin: a male pheromone that stimulates female memory and sexual attraction to an individual male's odour. BMC biology. 2010;8:75. Epub 2010/06/08. doi: 10.1186/1741-7007-8-75-. PubMed PMID: 20525243; PubMed Central PMCID: PMC2890510.

17. Roberts SA, Davidson AJ, McLean L, Beynon RJ, Hurst JL. Pheromonal induction of spatial learning in mice. Science. 2012;338(6113):1462–5. Epub 2012/12/15. doi: 10.1126/science.1225638. PubMed PMID: 23239735.

18. Armstrong SD, Robertson DH, Cheetham SA, Hurst JL, Beynon RJ. Structural and functional differences in isoforms of mouse major urinary proteins: a male-specific protein that preferentially binds a male pheromone. Biochem J. 2005;391(Pt 2):343–50. Epub 2005/06/07. doi: 10.1042/BJ20050404. PubMed PMID: 15934926; PubMed Central PMCID: PMC1276933.

19. Phelan MM, McLean L, Armstrong SD, Hurst JL, Beynon RJ, Lian LY. The structure, stability and pheromone binding of the male mouse protein sex pheromone darcin. PLoS One. 2014;9(10):e108415. doi: 10.1371/journal.pone.0108415. PubMed PMID: 25279835.

20. Berdoy M, Drickamer LC. Comparative social organization and life history of Rattus and Mus. Rodent societies: An ecological and evolutionary perspective Chicago, Illinois: University of Chicago Press; 2007. p. 380–92.

21. Vettorazzi A, Wait R, Nagy J, Monreal JI, Mantle P. Changes in male rat urinary protein profile during puberty: a pilot study. BMC Res Notes. 2013;6(1):232. Epub 2013/06/19. doi: 10.11861756-0500-6-232-. PubMed PMID: 23767887; PubMed Central PMCID: PMC3751546.

22. Guo X, Guo H, Zhao L, Zhang YH, Zhang JX. Two predominant MUPs, OBP3 and MUP13, are male pheromones in rats. Front Zool. 2018;15:6. doi: 10.1186/s12983-018-0254-0-. PubMed PMID: 29483934; PubMed Central PMCID: PMCPMC5824612.

23. Kumar V, Vasudevan A, Soh LJ, Le Min C, Vyas A, Zewail-Foote M, et al. Sexual attractiveness in male rats is associated with greater concentration of major urinary proteins. Biology of reproduction. 2014;91(6):150. doi: 10.1095/biolreprod.114.117903. PubMed PMID: 25359898.

24. Mudge JM, Armstrong SD, McLaren K, Beynon RJ, Hurst JL, Nicholson C, et al. Dynamic instability of the major urinary protein gene family revealed by genomic and phenotypic comparisons between C57 and 129 strain mice. Genome Biol. 2008;9(5):R91. Epub 2008/05/30. doi: 10.1186/gb-2008-9-5-r91. PubMed PMID: 18507838; PubMed Central PMCID: PMC2441477.

25. Sheehan MJ, Lee V, Corbett-Detig R, Bi K, Beynon RJ, Hurst JL, et al. Selection on Coding and Regulatory Variation Maintains Individuality in Major Urinary Protein Scent Marks in Wild Mice. Plos Genet. 2016;12(3):e1005891. doi: 10.1371/journal.pgen.1005891. PubMed PMID: 26938775; PubMed Central PMCID: PMCPMC4777540.

26. Logan DW, Marton TF, Stowers L. Species specificity in major urinary proteins by parallel evolution. PLoS One. 2008;3(9):e3280. Epub 2008/09/26. doi: 10.1371/journal.pone.0003280. PubMed PMID: 18815613; PubMed Central PMCID: PMC2533699.

27. Hancock JM. A bigger mouse? The rat genome unveiled. BioEssays : news and reviews in molecular, cellular and developmental biology. 2004;26(10):1039–42. Epub 2004/09/24. doi: 10.1002/bies.20121. PubMed PMID: 15382132.

28. Roy AK, Neuhaus OW. Identification of rat urinary proteins by zone and immunoelectrophoresis. Proceedings of the Society for Experimental Biology and Medicine Society for Experimental Biology and Medicine. 1966;121(3):894–9. Epub 1966/03/01. PubMed PMID: 4160706.

29. Kondo Y, Yamada J. Male urinary protein-1 (Mup-1) expression in the female rat. The Journal of heredity. 1983;74(4):280–2. Epub 1983/07/01. PubMed PMID: 6886376.

30. Vandoren G, Mertens B, Heyns W, Van Baelen H, Rombauts W, Verhoeven G. Different forms of alpha 2u-globulin in male and female rat urine. Eur J Biochem. 1983;134(1):175–81. Epub 1983/07/15. PubMed PMID: 6190651.

31. Aksu S, Tanrikulu F. Differentiation of protein species of alpha-2u-globulin according to database entries: A half-theoretical approach. J Proteomics. 2016;134:186–92. doi: 10.1016/j.jprot.2015.12.024. PubMed PMID: 26746007.

32. Lee RS, Monigatti F, Lutchman M, Patterson T, Budnik B, Steen JA, et al. Temporal variations of the postnatal rat urinary proteome as a reflection of systemic maturation. Proteomics. 2008;8(5):1097–112. Epub 2008/03/08. doi: 10.1002/pmic.200700701. PubMed PMID: 18324733.

33. Gibbs RA, Weinstock GM, Metzker ML, Muzny DM, Sodergren EJ, Scherer S, et al. Genome sequence of the Brown Norway rat yields insights into mammalian evolution. Nature. 2004;428(6982):493–521. doi: 10.1038/nature02426. PubMed PMID: 15057822.

34. Bayard C, Holmquist L, Vesterberg O. Purification and identification of allergenic alpha (2u)-globulin species of rat urine. Biochimica et biophysica acta. 1996;1290(2):129–34. Epub 1996/06/04. PubMed PMID: 8645715.

35. Bocskei Z, Groom CR, Flower DR, Wright CE, Phillips SE, Cavaggioni A, et al. Pheromone binding to two rodent urinary proteins revealed by X-ray crystallography. Nature. 1992;360(6400):186–8. Epub 1992/11/12. doi: 10.1038/360186a0. PubMed PMID: 1279439.

36. Chaudhuri BN, Kleywegt GJ, Bjorkman J, Lehman-McKeeman LD, Oliver JD, Jones TA. The structures of alpha 2u-globulin and its complex with a hyaline droplet inducer. Acta crystallographica Section D, Biological crystallography. 1999;55(Pt 4):753–62. Epub 1999/03/25. PubMed PMID: 10089305.

37. Gómez-Baena G, Armstrong SD, Phelan MM, Hurst JL, Beynon RJ. The major urinary protein system in the rat. Biochem Soc T. 2014;42:886–92. doi: 10.1042/Bst20140083. PubMed PMID: WOS:000340329200027.

38. Knopf JL, Gallagher JF, Held WA. Differential, multihormonal regulation of the mouse major urinary protein gene family in the liver. Mol Cell Biol. 1983;3(12):2232–40. Epub 1983/12/01. PubMed PMID: 6656765; PubMed Central PMCID: PMC370094.

39. Kuhn NJ, Woodworth-Gutai M, Gross KW, Held WA. Subfamilies of the mouse major urinary protein (MUP) multi-gene family: sequence analysis of cDNA clones and differential regulation in the liver. Nucleic Acids Res. 1984;12(15):6073–90. PubMed PMID: 6548015; PubMed Central PMCID: PMCPMC320058.

40. Kulkarni AB, Gubits RM, Feigelson P. Developmental and hormonal regulation of alpha 2u-globulin gene transcription. Proc Natl Acad Sci U S A. 1985;82(9):2579–82. Epub 1985/05/01. PubMed PMID: 2581250; PubMed Central PMCID: PMC397607.

41. MacInnes JI, Nozik ES, Kurtz DT. Tissue-specific expression of the rat alpha 2u globulin gene family. Mol Cell Biol. 1986;6(10):3563–7. Epub 1986/10/01. PubMed PMID: 2432391; PubMed Central PMCID: PMC367109.

42. Murty CV, Mancini MA, Chatterjee B, Roy AK. Changes in transcriptional activity and matrix association of alpha 2u-globulin gene family in the rat liver during maturation and aging. Biochim Biophys Acta. 1988;949(1):27–34. Epub 1988/01/25. PubMed PMID: 2446666.

43. Saito K, Nishikawa J, Imagawa M, Nishihara T, Matsuo M. Molecular evidence of complex tissue- and sex-specific mRNA expression of the rat alpha(2u)-globulin multigene family. Biochem Biophys Res Commun. 2000;272(2):337–44. Epub 2000/06/02. doi: 10.1006/bbrc.2000.2694. PubMed PMID: 10833415.

44. Elliott BM, Ramasamy R, Stonard MD, Spragg SP. Electrophoretic variants of alpha 2u-globulin in the livers of adult male rats: a possible polymorphism. Biochim Biophys Acta. 1986;870(1):135–40. Epub 1986/03/07. PubMed PMID: 2418881.

45. Wait R, Gianazza E, Eberini I, Sironi L, Dunn MJ, Gemeiner M, et al. Proteins of rat serum, urine, and cerebrospinal fluid: VI. Further protein identifications and interstrain comparison. Electrophoresis. 2001;22(14):3043–52. Epub 2001/09/22. doi: 10.1002/1522-2683(200108)22:14<3043::AID-ELPS3043>3.0.CO;2-M. PubMed PMID: 11565799.

46. Payne CE, Malone N, Humphries RE, Bradbook C, Veggerby C, Beynon RJ, et al. Heterogeneity of major urinary proteins in the house mice: population and sex differences. In: Marchelewska-Koj A, Muller-Schwarze D, Lepri J, editors. Chemical Signals in Vertebrates. New York: Plenum Press; 2001. p. 233–40.

47. Beynon RJ, Veggerby C, Payne CE, Robertson DH, Gaskell SJ, Humphries RE, et al. Polymorphism in major urinary proteins: molecular heterogeneity in a wild mouse population. Journal of chemical ecology. 2002;28(7):1429–46. Epub 2002/08/30. PubMed PMID: 12199505.

48. Thom MD, Stockley P, Jury F, Ollier WE, Beynon RJ, Hurst JL. The direct assessment of genetic heterozygosity through scent in the mouse. Current biology : CB. 2008;18(8):619–23. Epub 2008/04/22. doi: 10.1016/j.cub.2008.03.056. PubMed PMID: 18424142.

49. Sherborne AL, Thom MD, Paterson S, Jury F, Ollier WE, Stockley P, et al. The genetic basis of inbreeding avoidance in house mice. Current biology : CB. 2007;17(23):2061–6. Epub 2007/11/13. doi: 10.1016/j.cub.2007.10.041. PubMed PMID: 17997307; PubMed Central PMCID: PMC2148465.

50. Drickamer K, Kwoh TJ, Kurtz DT. Amino acid sequence of the precursor of rat liver alpha 2 micro-globulin. The Journal of biological chemistry. 1981;256(8):3634–6. Epub 1981/04/25. PubMed PMID: 6163771.

51. Beynon RJ, Armstrong SD, Claydon AJ, Davidson AJ, Eyers CE, Langridge JI, et al. Mass spectrometry for structural analysis and quantification of the Major Urinary Proteins of the house mouse. Int J Mass Spectrom. 2015;391:146–56. doi: 10.1016/j.ijms.2015.07.026. PubMed PMID: WOS:000367124800018.

52. Mertens B, Verhoeven G. Influence of neonatal androgenization on the expression of alpha 2u-globulin in rat liver and submaxillary gland. J Steroid Biochem. 1985;23(5A):557–65. Epub 1985/11/01. PubMed PMID: 2417039.

53. Papes F, Logan DW, Stowers L. The vomeronasal organ mediates interspecies defensive behaviors through detection of protein pheromone homologs. Cell. 2010;141(4):692–703. Epub 2010/05/19. doi: 10.1016/j.cell.2010.03.037. PubMed PMID: 20478258; PubMed Central PMCID: PMC2873972.

54. Ichiyoshi Y, Endo H, Yamamoto M. Length polymorphism in the 3' noncoding region of rat hepatic alpha 2u-globulin mRNAs. Biochimica et biophysica acta. 1987;910(1):43–51. Epub 1987/10/09. PubMed PMID: 2443176.

55. Gao F, Endo H, Yamamoto M. Length heterogeneity in rat salivary gland alpha 2 mu globulin mRNAs: multiple splice-acceptors and polyadenylation sites. Nucleic Acids Res. 1989;17(12):4629–36. Epub 1989/06/26. PubMed PMID: 2473438; PubMed Central PMCID: PMC318020.

56. Lobel D, Strotmann J, Jacob M, Breer H. Identification of a third rat odorant-binding protein (OBP3). Chemical senses. 2001;26(6):673–80. Epub 2001/07/28. PubMed PMID: 11473933.

57. Saito H, Chi Q, Zhuang H, Matsunami H, Mainland JD. Odor coding by a Mammalian receptor repertoire. Science signaling. 2009;2(60):ra9. Epub 2009/03/06. doi: 10.1126/scisignal.2000016. PubMed PMID: 19261596; PubMed Central PMCID: PMC2774247.

58. Laperche Y, Lynch KR, Dolan KP, Feigelson P. Tissue-specific control of alpha 2u globulin gene expression: constitutive synthesis in the submaxillary gland. Cell. 1983;32(2):453–60. Epub 1983/02/01. PubMed PMID: 6186396.

59. Tagliabracci VS, Wiley SE, Guo X, Kinch LN, Durrant E, Wen J, et al. A Single Kinase Generates the Majority of the Secreted Phosphoproteome. Cell. 2015;161(7):1619–32. doi: 10.1016/j.cell.2015.05.028. PubMed PMID: 26091039; PubMed Central PMCID: PMCPMC4963185.

60. Rajkumar R, Ilayaraja R, Alagendran S, Archunan G, Maralidharan AR, Huang PH, et al. Characterization of rat odorant binding protein variants and its post-translational modifications (PTMs):LC-MS/MS analyses of protein Eluted from 2D-Polyacrylamide gel electrophoresis. Proteomics and Bioinformatics. 2011;4(10):210–7.

61. Brimau F, Cornard JP, Le Danvic C, Lagant P, Vergoten G, Grebert D, et al. Binding specificity of recombinant odorant-binding protein isoforms is driven by phosphorylation. Journal of chemical ecology. 2010;36(8):801–13. doi: 10.1007/s10886-010-9820-4. PubMed PMID: 20589419.

62. Nielsen H. Predicting Secretory Proteins with SignalP. Methods Mol Biol. 2017;1611:59–73. doi: 10.1007/978-1-4939-7015-5_6. PubMed PMID: 28451972.

63. Beynon RJ, Oliver S, Robertson DH. Characterization of the soluble, secreted form of urinary meprin. Biochem J. 1996;315 (Pt 2):461–5. PubMed PMID: 8615815; PubMed Central PMCID: PMCPMC1217218.

64. Laemmli UK. Cleavage of structural proteins during the assembly of the head of bacteriophage T4. Nature. 1970;227(5259):680–5. PubMed PMID: 5432063.

65. Hammond DE, Claydon AJ, Simpson DM, Edward D, Stockley P, Hurst JL, et al. Proteome Dynamics: Tissue Variation in the Kinetics of Proteostasis in Intact Animals. Mol Cell Proteomics. 2016;15(4):1204–19. doi: 10.1074/mcp.M115.053488. PubMed PMID: 26839000; PubMed Central PMCID: PMCPMC4824850.

66. Beynon RJ. A simple tool for drawing proteolytic peptide maps. Bioinformatics. 2005;21(5):674–5. PubMed PMID: 15539446.

67. Strausberg RL, Feingold EA, Grouse LH, Derge JG, Klausner RD, Collins FS, et al. Generation and initial analysis of more than 15,000 full-length human and mouse cDNA sequences. Proc Natl Acad Sci U S A. 2002;99(26):16899–903. doi: 10.1073/pnas.242603899. PubMed PMID: 12477932; PubMed Central PMCID: PMCPMC139241.

68. Shannon P, Markiel A, Ozier O, Baliga NS, Wang JT, Ramage D, et al. Cytoscape: a software environment for integrated models of biomolecular interaction networks. Genome Res. 2003;13(11):2498–504. doi: 10.1101/gr.1239303. PubMed PMID: 14597658; PubMed Central PMCID: PMCPMC403769.

