## Supplementary figures and images for "Molecular complexity of the major urinary protein system of the Norway rat, *Rattus norvegicus*"

### Supplementary file 1

Supplementary Fig 1

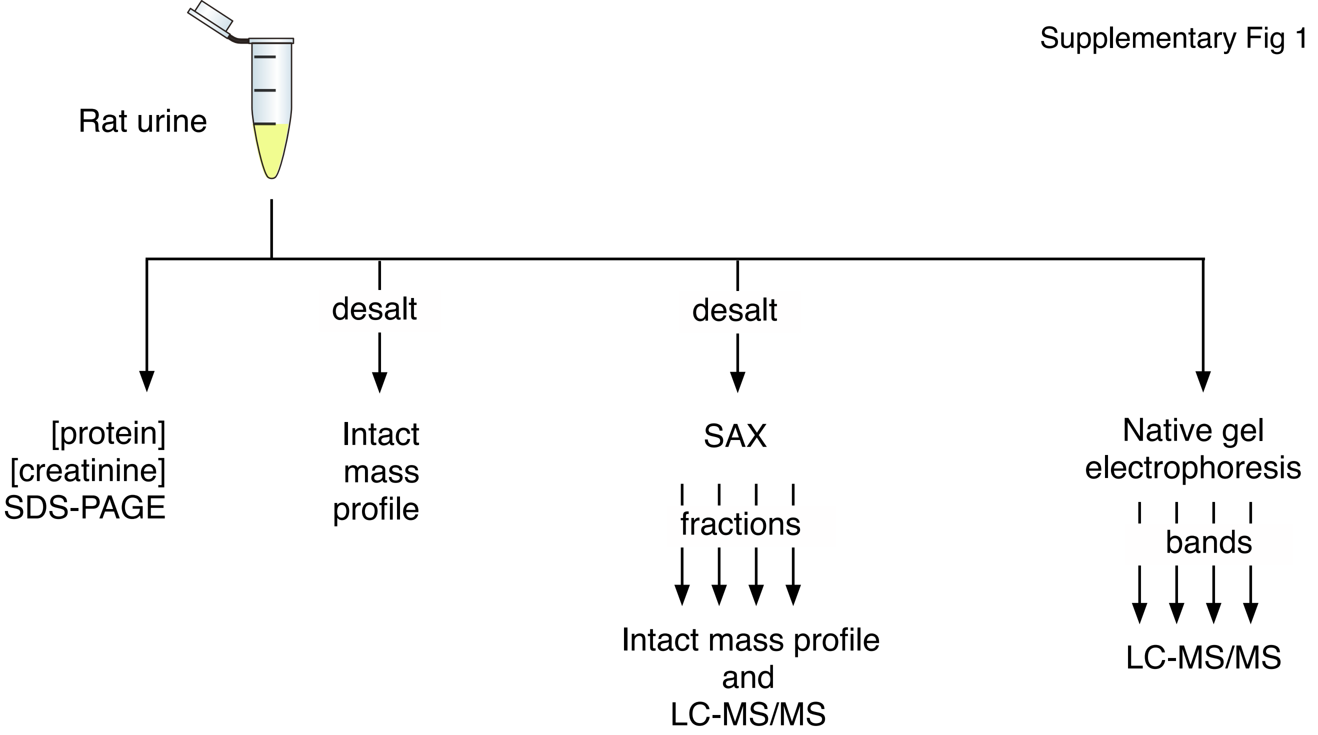

### Supplementary file 4

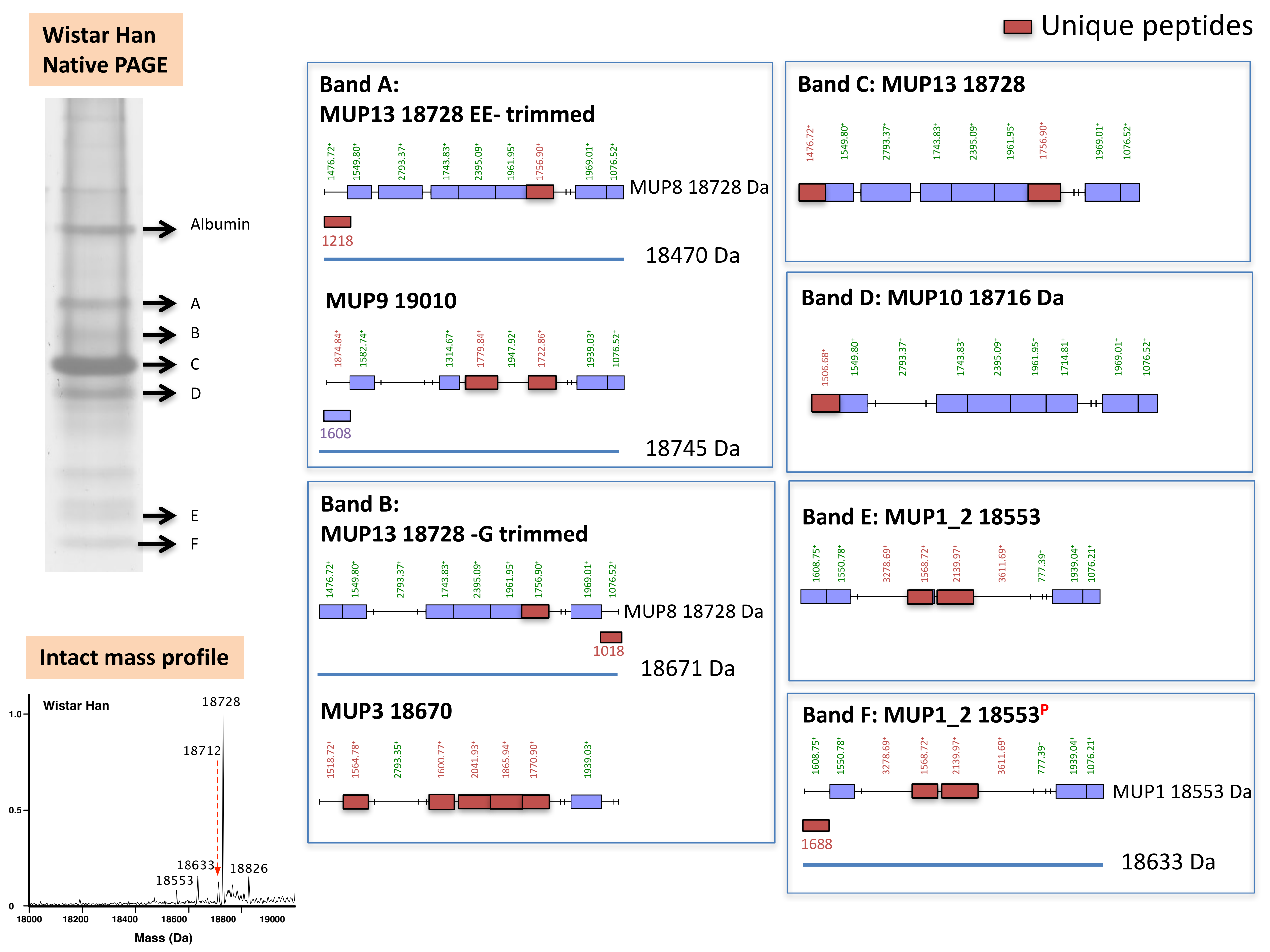

### Supplementary file 5

##### Brown Norway Native PAGE

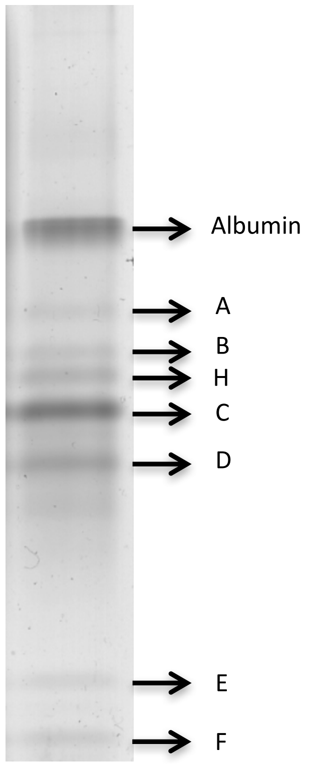


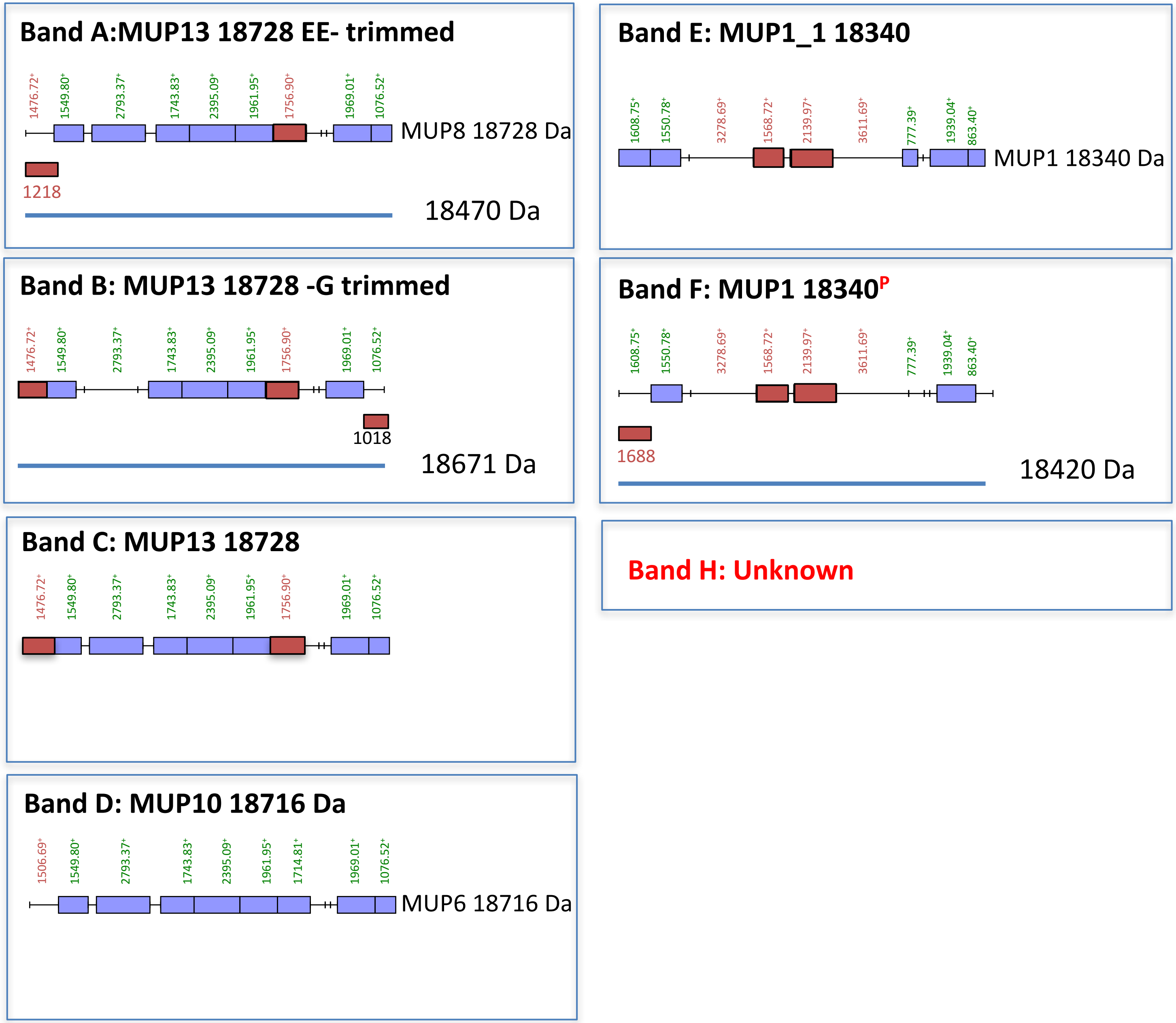


##### Intact mass profile

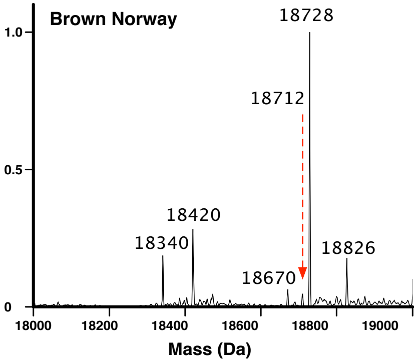

### Supplementary file 6

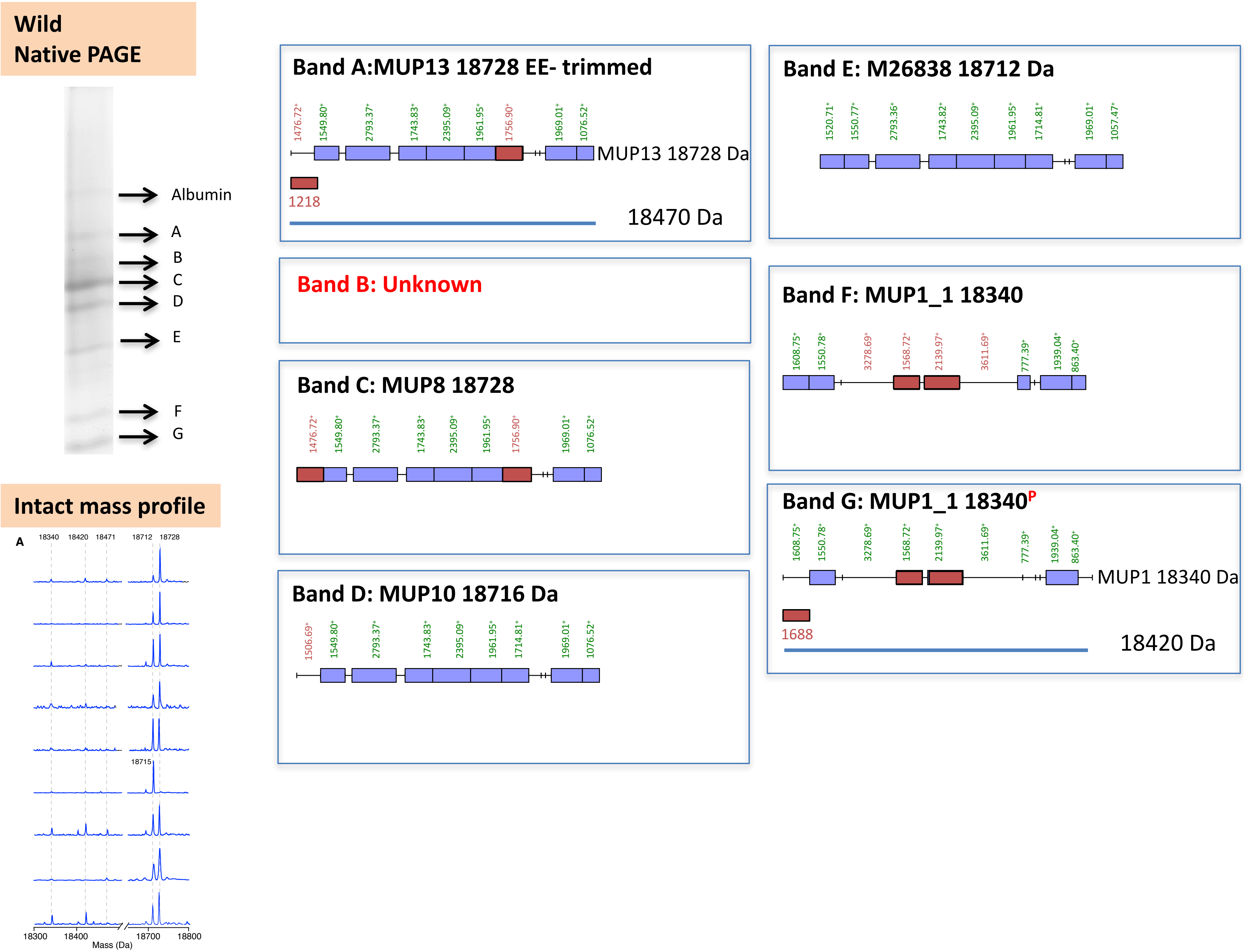

### Supplementary file 7

#### Wistar Han male: SAX

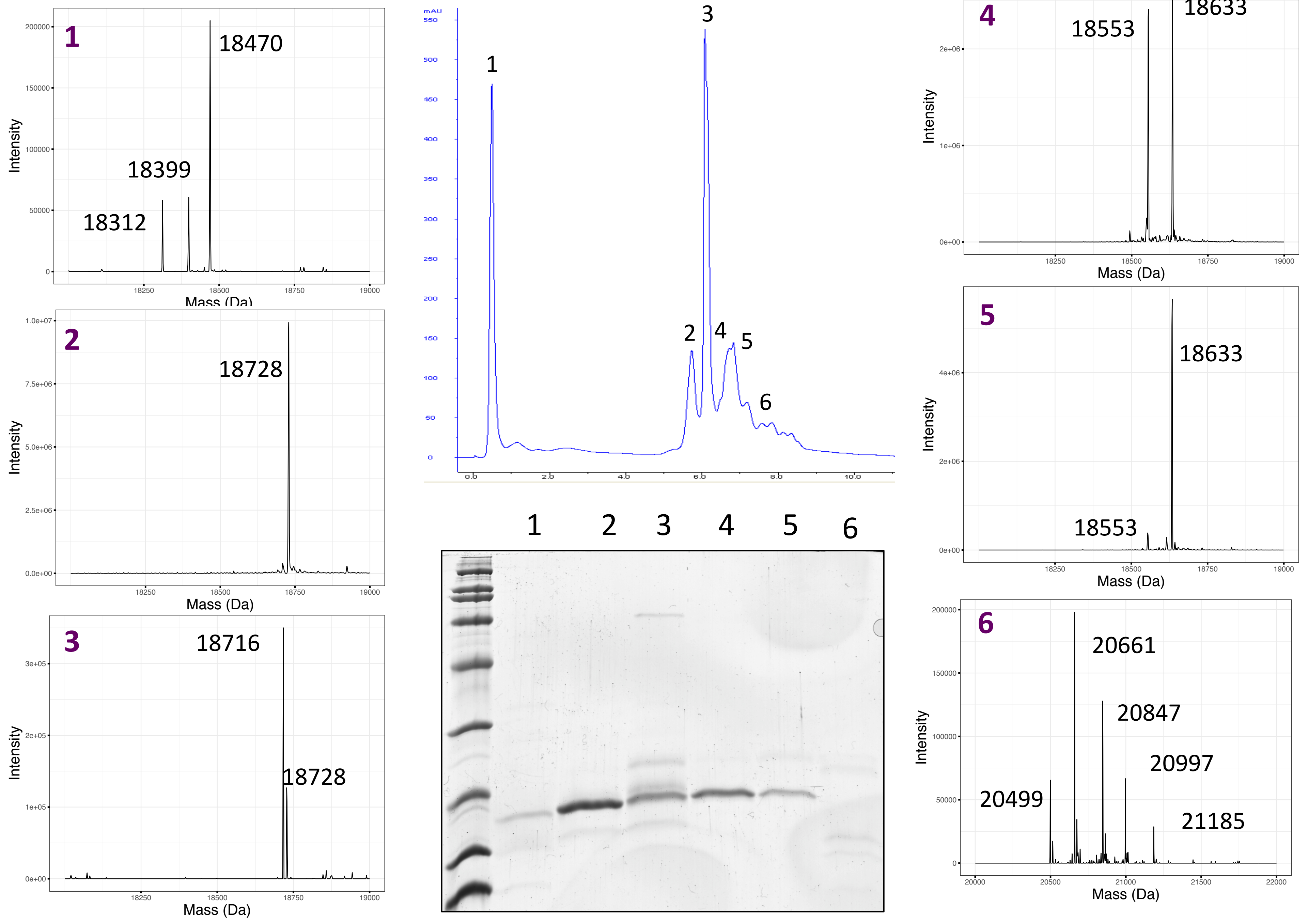

### Supplementary file 8

#### Brown Norway male: ion exchange chromatography

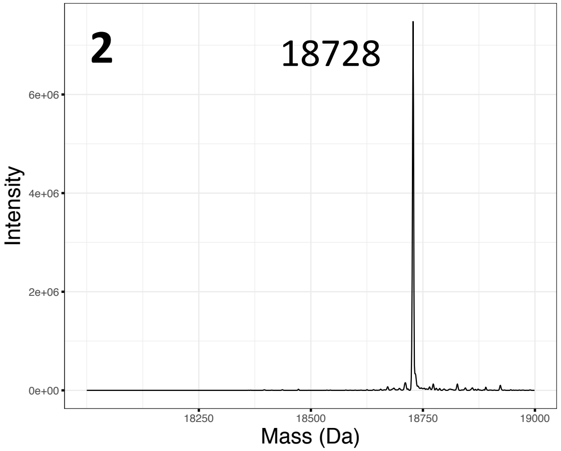


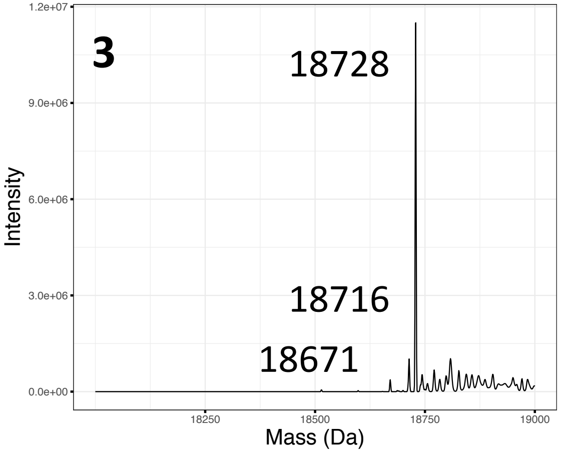


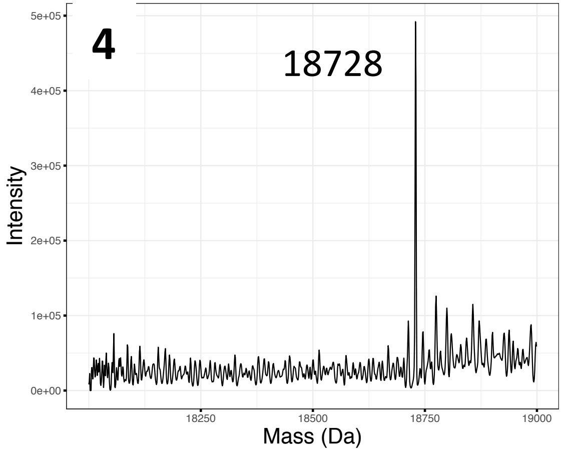


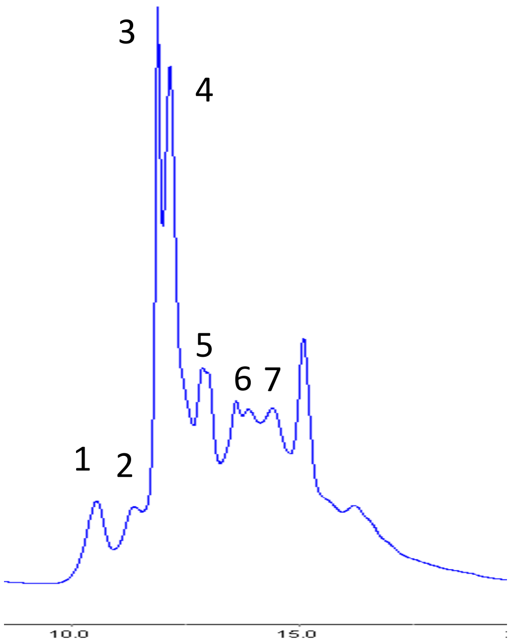


*No good spectra for fraction 1 and 5

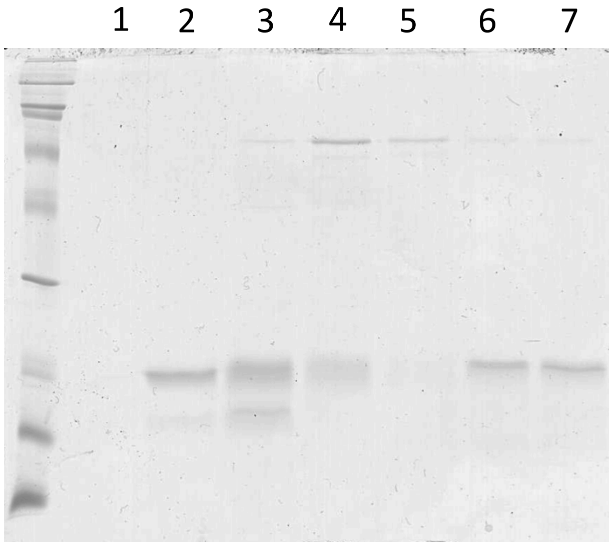


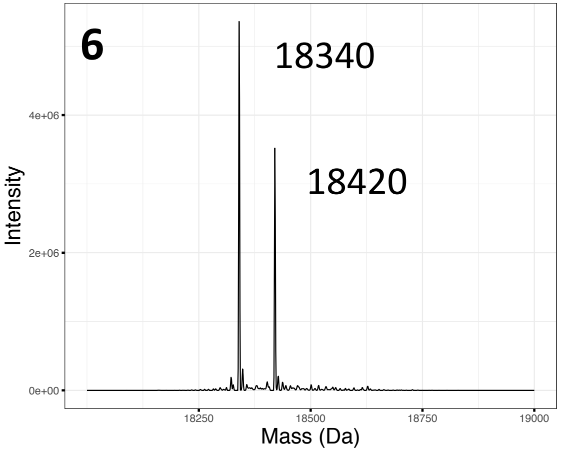


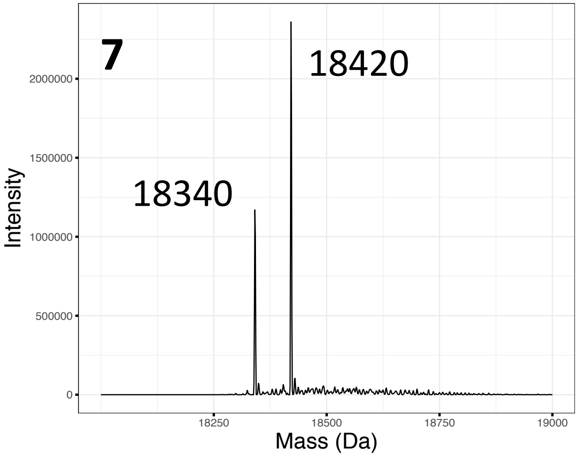

### Supplementary file 9

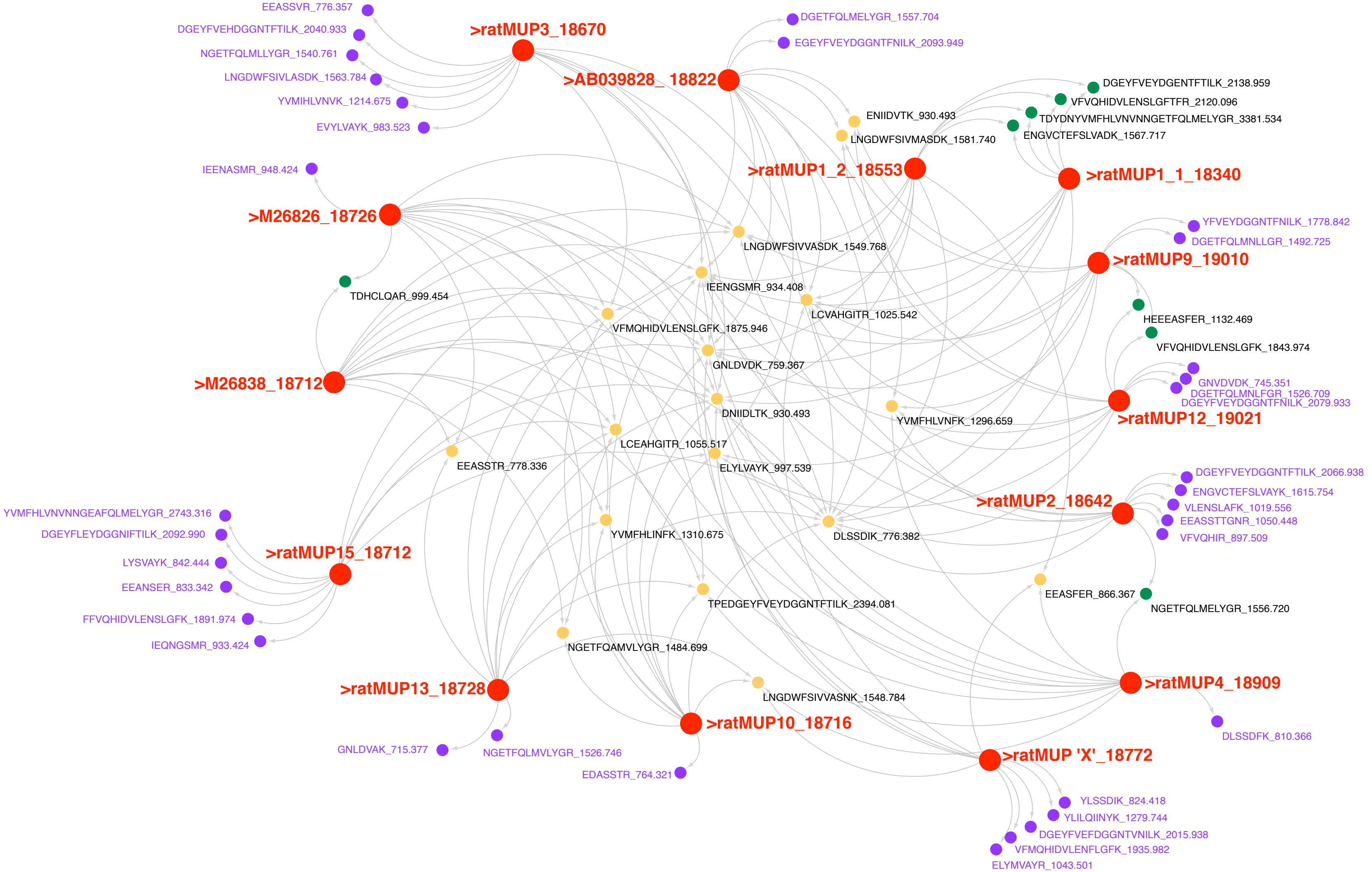
