## Supplementary material for "Molecular complexity of the major urinary protein system of the Norway rat, *Rattus norvegicus*"

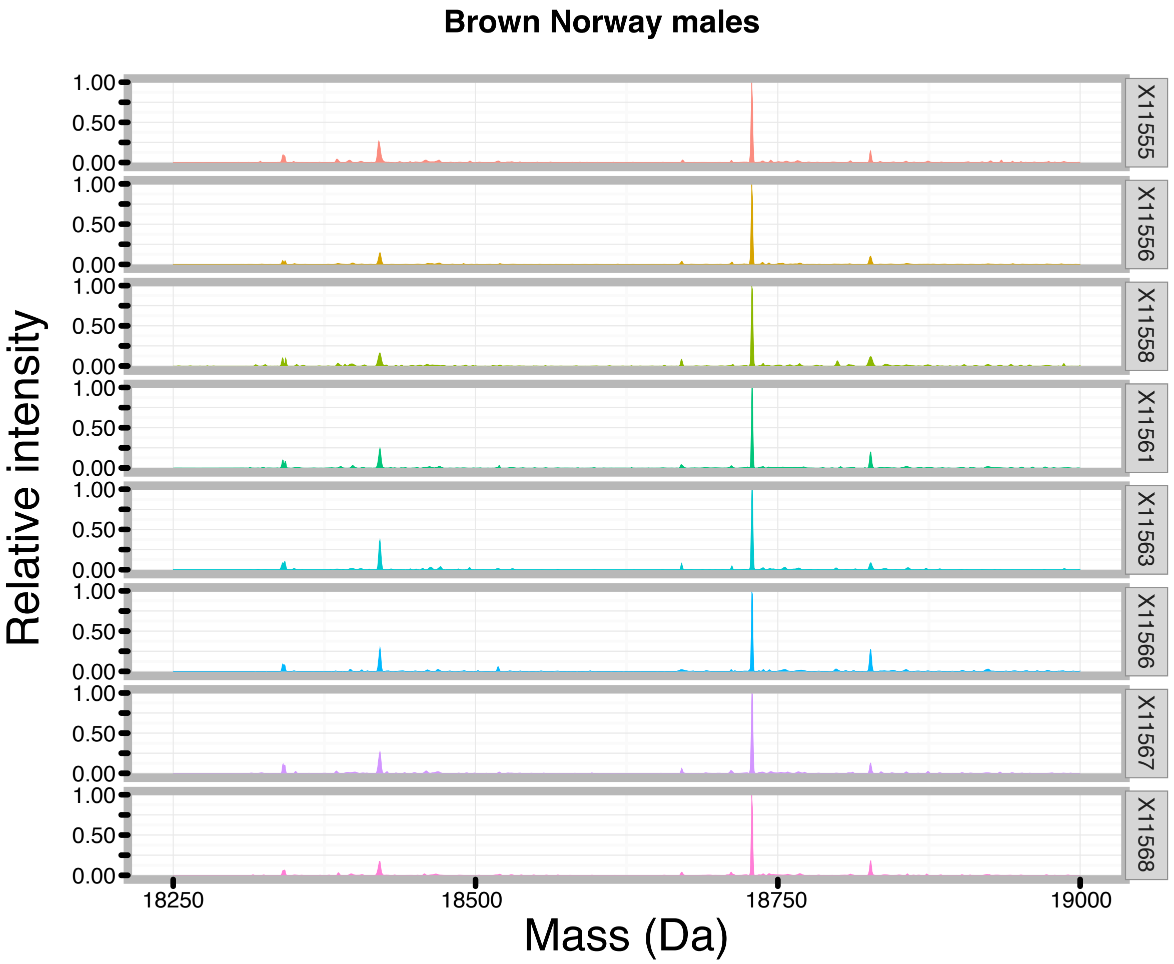


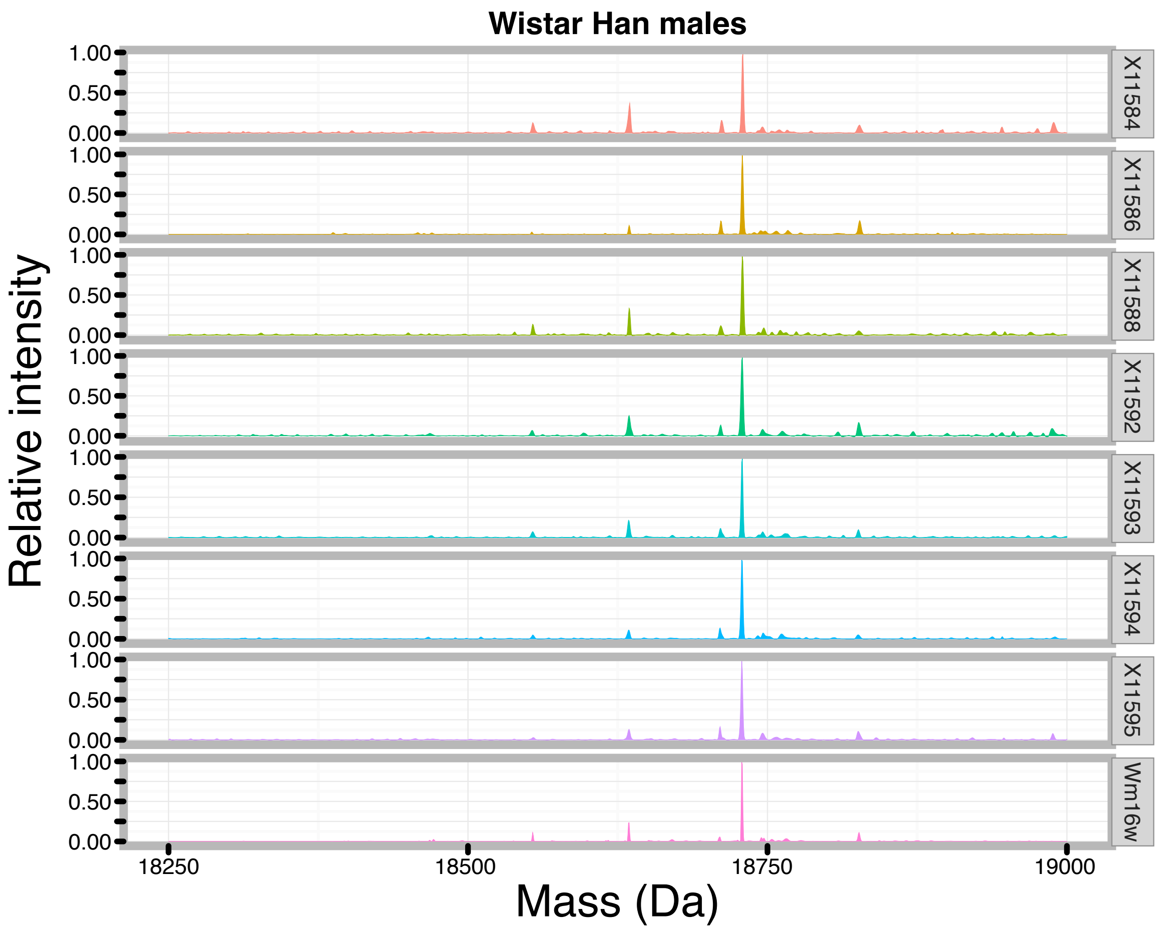


Supplementary Figure 2: Individual intact mass spectra from Wistar Han and Brown Norway animals.
