## Supplementary material for "Molecular complexity of the major urinary protein system of the Norway rat, *Rattus norvegicus*"

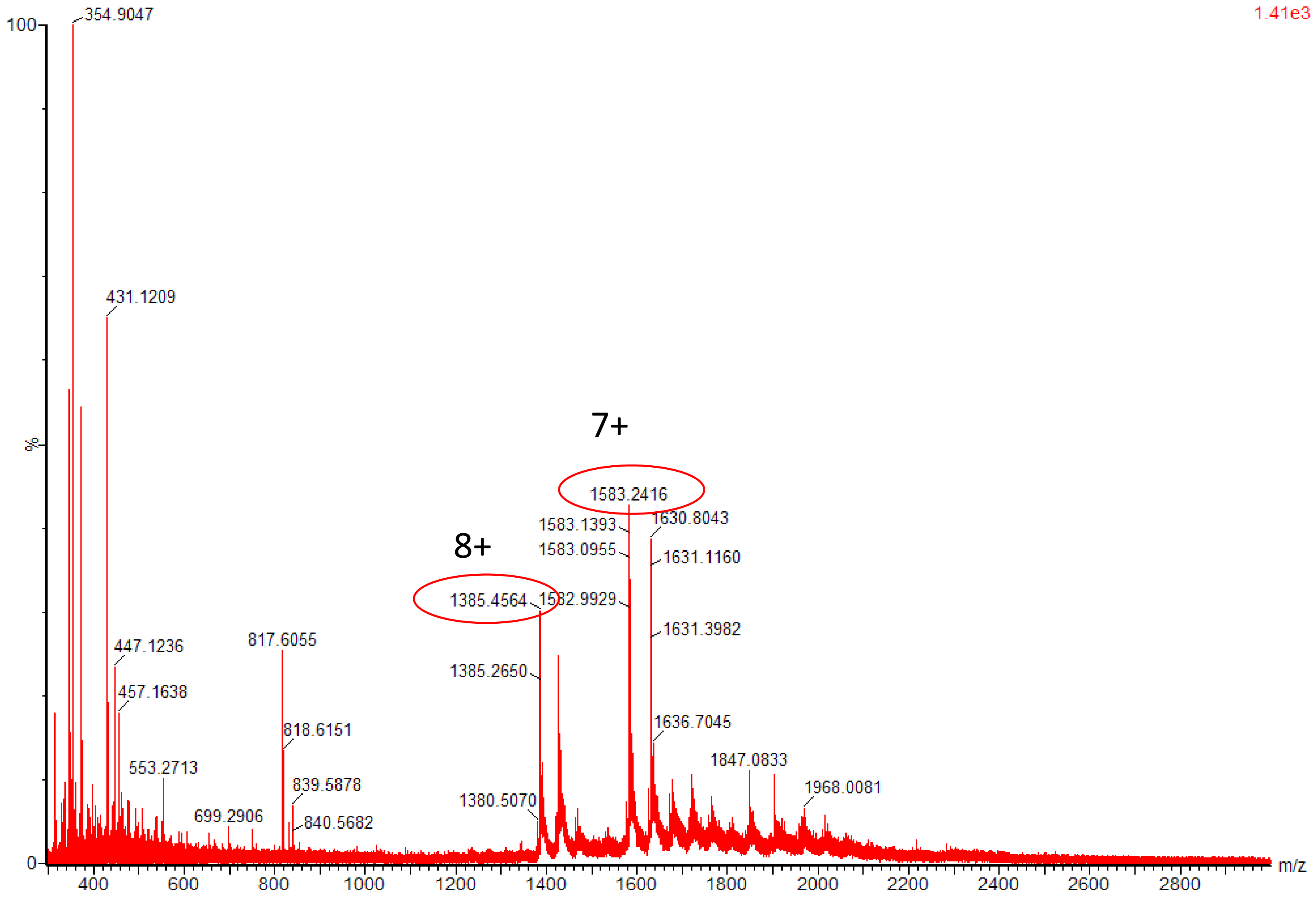


Supplementary Figure 3: ESI-MS analysis of female rat urine. Deconvolution of the spectrum estimates two masses of about 11 kDa (11065 and 11450 Da) likely corresponding to the rat urinary proteins 1 and 2.
